# Distinguishing between recruitment and spread of silent chromatin structures in *Saccharomyces cerevisiae*

**DOI:** 10.1101/2021.11.25.469980

**Authors:** Molly Brothers, Jasper Rine

## Abstract

The formation of heterochromatin at *HML*, *HMR*, and telomeres in *Saccharomyces cerevisiae* involves two main steps: Recruitment of Sir proteins to silencers and their spread throughout the silenced domain. We developed a method to study these two processes at single base-pair resolution. Using a fusion protein between the heterochromatin protein Sir3 and the non-site-specific bacterial adenine methyltransferase M.EcoGII, we mapped sites of Sir3-chromatin interactions genome-wide using long-read Nanopore sequencing to detect adenines methylated by the fusion protein. A silencing-deficient mutant of Sir3 lacking its Bromo-Adjacent Homology (BAH) domain, *sir3-bahΔ*, was still recruited to *HML*, *HMR*, and telomeres. However, in the absence of the BAH domain, it was unable to spread away from those recruitment sites. Overexpression of Sir3 did not lead to further spreading at *HML*, *HMR*, and most telomeres. A few exceptional telomeres, like 6R, exhibited a small amount of Sir3 spreading, suggesting that boundaries at telomeres responded variably to Sir3 overexpression. Finally, by using a temperature-sensitive allele of *SIR3* fused to *M.ECOGII*, we tracked the positions first methylated after induction and found that repression of genes at *HML* and *HMR* began before Sir3 occupied the entire locus.

## INTRODUCTION

Cells have an interest in coordinating the expression of genes: It allows them to turn sets of genes on and off in response to various stimuli or ensure certain genes are always expressed or always repressed to create and maintain cell identity. There are multiple ways to coordinate transcription, including shared binding sites for activators or repressors in promoters, nuclear compartmentalization, and creation of large domains like heterochromatin. The establishment of coordinated, stable blocs of gene expression, such as heterochromatin, can be broken down into two main steps: Nucleation, which involves the recruitment of chromatin-modifying factors, followed by the expansion of these chromatin-modifying factors beyond recruitment sites in an ill-defined process known as spreading.

An impressive example of the concepts of nucleation and spread is inactivation of the X chromosome in female mammals (reviewed in Galupa and Heard, 2018). Nucleation begins with the transcription of the non-coding RNA *Xist* from one of the two X chromosomes, which recruits heterochromatin factors in *cis* that eventually coat and transcriptionally silence nearly the entire 167-megabase X chromosome. Two studies have characterized early steps in nucleation at the *Xist* locus and at recruitment sites throughout the X chromosome (Engreitz et al., 2013; Simon et al., 2013), but the mechanics of how the XIST transcript and associated heterochromatin proteins spread remains unclear even for this well-studied phenomenon. Furthermore, ‘spreading’ itself is an inferred process that connects known recruitment sites to the final binding profile of heterochromatin proteins. A clear mechanistic distinction between nucleation and spread, with a characterization of intermediate steps, has not been achieved for any organism.

Transcriptional silencing in the budding yeast *Saccharomyces cerevisiae* is one of the best-studied heterochromatic phenomena (reviewed in Gartenberg and Smith, 2016). Heterochromatin is created by the <u>S</u>ilent Information <u>R</u>egulator (SIR) complex that silences the transcription of genes at *HML* and *HMR* and the 32 telomeres. Recruitment of the SIR complex to *HML*, *HMR*, and telomeres is sequence specific, whereas its spreading is sequence nonspecific. More specifically, different combinations of Rap1, Abf1, and Origin Recognition Complex (ORC) binding sites are present at the *E* and *I* silencers that flank *HML* and *HMR* (Buchman et al., 1988; Kimmerly et al., 1988; Shore et al., 1987; Shore and Nasmyth, 1987) and at the TG repeats and X elements of telomeres (Buchman et al., 1988; Longtine et al., 1989; Stavenhagen and Zakian, 1994). These proteins in turn interact with and recruit the SIR complex (Cockell et al., 1995; Moretti and Shore, 2001; Triolo and Sternglanz, 1996). The SIR complex then deacetylates chromatin (Braunstein et al., 1993; Ellahi et al., 2015; Suka et al., 2001; Thurtle and Rine, 2014), resulting in chromatin compaction (Georgel et al., 2001; Gottschling, 1992; Johnson et al., 2009; Loo and Rine, 1994; Singh and Klar, 1992; Swygert et al., 2018). As a result, transcription is blocked at least in part by steric occlusion, though details remain unknown (Chen and Widom, 2005; Gao and Gross, 2008; Johnson et al., 2013; Lynch and Rusche, 2009; Sekinger and Gross, 2001; Steakley and Rine, 2015). Almost any gene placed within the defined domain can be transcriptionally silenced, establishing the locus-specific, gene non-specific nature of heterochromatic silencing (Dodson and Rine, 2015; Gottschling et al., 1990; Saxton and Rine, 2019; Schnell and Rine, 1986; Sussel et al., 1993). This difference in sequence dependence between recruitment and spread implies they are separable processes that rely on different factors and interactions.

Of the three SIR complex members (Sir2, Sir3, and Sir4), Sir3 is thought to be the major structural driver of heterochromatin spread and compaction. Sir3 interacts with the silencer-binding proteins Abf1 and Rap1 (Moretti et al., 1994; Moretti and Shore, 2001), with the other members of the SIR complex, Sir2 and Sir4 (Chang et al., 2003; Ehrentraut et al., 2011; Rudner et al., 2005; Samel et al., 2017; Strahl-Bolsinger et al., 1997), with nucleosomes (Armache et al., 2011; Hecht et al., 1995; Johnson et al., 1990; Norris et al., 2008; Onishi et al., 2007), and with itself (King et al., 2006; Liaw and Lustig, 2006; Oppikofer et al., 2013). All of these Sir3 interactions are required for transcriptional silencing. *In vitro*, Sir3 dimers can bridge neighboring nucleosomes and compact chromatin (Behrouzi et al., 2016). Among the SIR complex members, Sir3 has the largest difference in affinity for acetylated and deacetylated histone tails (Armache et al., 2011; Carmen et al., 2002; Onishi et al., 2007; Oppikofer et al., 2011; Swygert et al., 2018). By binding deacetylated histone tails more strongly than acetylated ones, Sir3 helps create a positive feedback loop wherein Sir2 deacetylates histone tails (Ghidelli et al., 2001; Imai et al., 2000; Landry et al., 2000; Smith et al., 2000) and Sir3 reinforces and further recruits Sir2/4 to silent regions.

Characterization of SIR complex nucleation and spread is limited by techniques like ChIP-seq that measure processes on populations of molecules and at a resolution limited by sequencing-read length. We developed a new method that allowed us to characterize the binding of heterochromatin proteins at base-pair resolution. Using long-read sequencing, we used this new method to resolve the distinction between the recruitment and spread of Sir3 and to track the establishment of heterochromatin over time. These data pinpointed when the process of transcriptional silencing begins.

## RESULTS

### The Sir3-M.EcoGII fusion protein strongly and specifically methylated *HML* and *HMR*

To achieve a higher-resolution method for assessing SIR complex binding, we made a fusion protein between Sir3 and M.EcoGII (Figure 1A), a non-site-specific bacterial N6-methyladenosine methyltransferase (Murray et al., 2018; Woodcock et al., 2020). In principle, wherever Sir3 binds chromatin, even transiently, M.EcoGII should methylate nearby accessible adenines to make m^6^A. *S. cerevisiae* has no endogenous DNA methylation and no demethylases, allowing us to attribute m^6^A only to the activity of the fusion protein, reflecting where it resides as well as where it has been. M. EcoGII has no specific recognition sequencing, which should provide more resolution than methyltransferases like *E. coli* Dam, which has a four base-pair recognition site. The positions of Sir3-M.EcoGII can be determined conventionally by immunoprecipitation with an antibody against m^6^A followed by sequencing the precipitated DNA using Illumina sequencing. The more powerful implementation would come from distinguishing between individual m^6^A bases and unmodified A bases using long-read Nanopore sequencing. (Xu and Seki, 2020).

**Figure 1.**
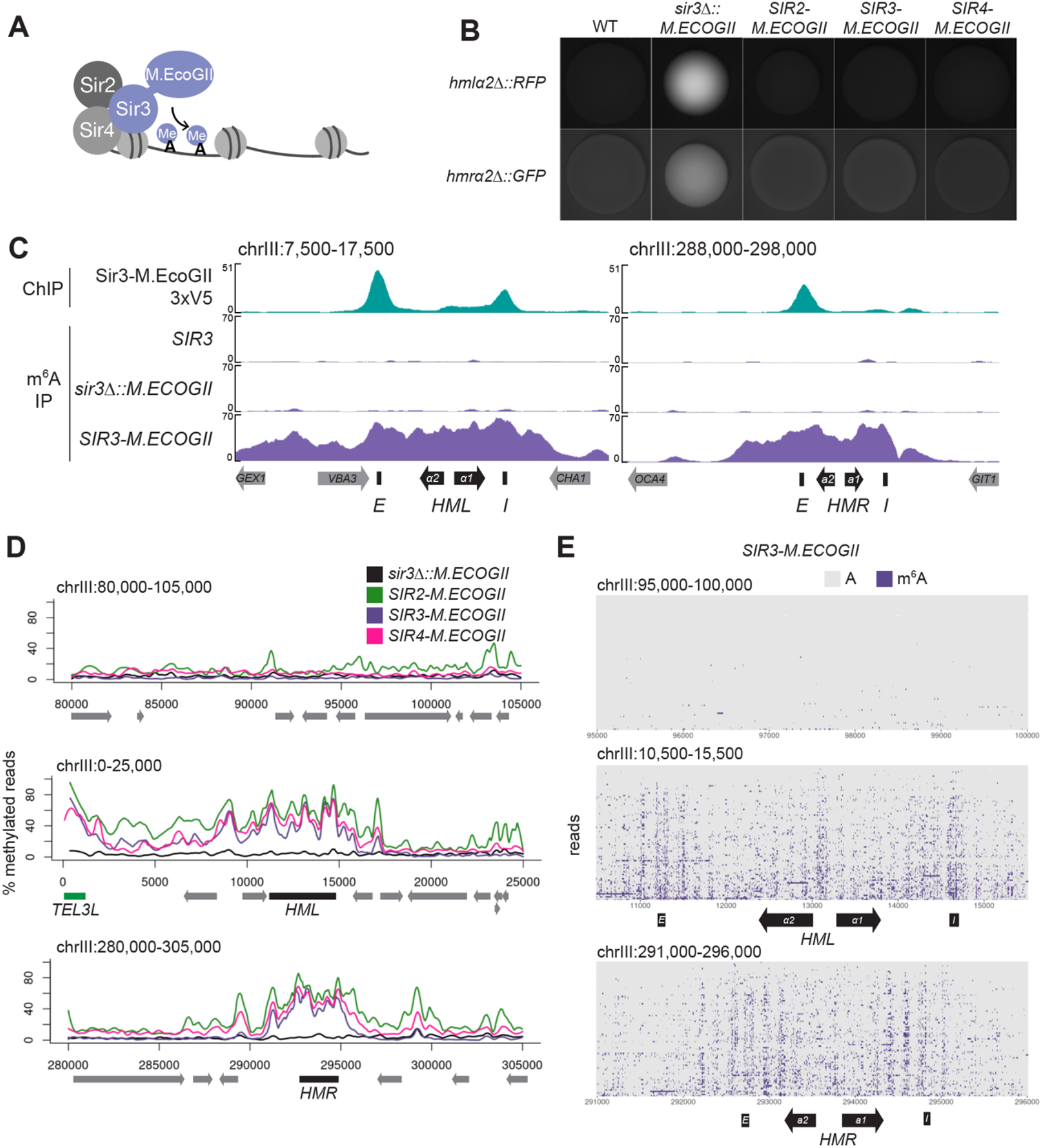
The Sir3-M.EcoGII fusion protein strongly and specifically methylated *HML* and *HMR*. **A)** Sir3-M.EcoGII is a fusion protein that non-specifically methylates adenines in regions that Sir3 binds. **B)** Genes encoding fluorescence reporters were placed at *HMLα2* (*RFP*) and *HMRα2* (*GFP*) to report on transcription from the two loci. Shown are representative images of colonies from each strain: no M.EcoGII (WT, JRY12731), unfused M.EcoGII (*sir3Δ::M.EcoGII*, JRY12842), *SIR2-M.ECOGII* (JRY13660), *SIR3-M.ECOGII* (JRY12844), and *SIR4-M.ECOGII* (JRY13019). **C)** ChIP-seq of Sir3-M.EcoGII-3xV5 (top row, JRY12839) and DNA m6A immunoprecipitation and sequencing (DIP-seq) of no EcoGII (row two, JRY11699), *sir3Δ::M.ECOGII* (row three, JRY12838), and *SIR3-M.ECOGII* (row four, JRY12840). Shown are 10 kb regions centered at *HML* (left) and *HMR* (right). Input results are plotted but not visible due to the strong ChIP-seq and DIP-seq signals. **D)** Aggregate results from long-read Nanopore sequencing of *sir3Δ::M.ECOGII* (black line, JRY12838), *SIR2-M.ECOGII* (green line, JRY13625), *SIR3-M.ECOGII* (purple line, JRY13027), and *SIR4-M.ECOGII* (pink line, JRY13021). The y-axis represents the percentage of reads in each position called as methylated by the modified-base calling software Megalodon (see Materials & Methods). Shown are 25 kb windows at a control region on chromosome III to show background methylation (top row), at *HML* (middle row), and at *HMR* (bottom row). **E)** Single-read plots from long-read Nanopore sequencing of *SIR3-M.ECOGII* (JRY13027). Each row of the plots is a single read the spans the entire query region, ordered by lowest average methylation on the top to highest average methylation on the bottom. Methylated adenines are colored purple, and unmethylated adenines are colored gray. Shown are 5 kb windows at a control region on chromosome III to show background methylation (top row), at *HML* (middle row), and at *HMR* (bottom row)

To test this concept, we first assessed the silencing ability of Sir3-M.EcoGII by growing colonies with GFP integrated at *HMRα2* and RFP integrated at *HMLα2*. Colonies expressing Sir3-M.EcoGII produced colonies with no fluorescence (Figure 1B). Thus, the fusion protein retained full function of Sir3.

To compare the binding profile of Sir3-M.EcoGII to the distribution of methylation it produced, we performed DNA immunoprecipitation and Illumina sequencing (DIP-seq) using an antibody that specifically recognizes m^6^A alongside ChIP-seq for a V5 epitope-tagged Sir3-M.EcoGII. ChIP-seq revealed strong Sir3-M.EcoGII occupancy over the *E* and *I* silencers at *HML* and over the *E* silencer at *HMR* with weaker but consistent signal above background between the two silencers (Figure 1C, top row). Compared to ChIP-seq, methylation measured by DIP-seq had a stronger signal over the entirety of *HML* and *HMR* and a broader signal that extended beyond the silencers (Figure 1C, Figure 1–figure supplement 1). The methylation over *HML* and *HMR* was from the fusion protein Sir3-M.EcoGII, as neither a strain without M.EcoGII nor a strain expressing unfused M.EcoGII from the *SIR3* promoter showed appreciable DIP-seq signal (Figure 1C, Figure 1–figure supplement 1).

Methylation by Sir3-M.EcoGII measured by Nanopore sequencing agreed well with DIP-seq, showing strong methylation over *HML* and *HMR* with little background methylation outside of these regions (Figure 1D, Figure 1–figure supplement 2A). Fusions of Sir2 and Sir4 with M.EcoGII were also fully silencing competent (Figure 1B) and produced methylation signals that matched Sir3-M.EcoGII at *HML* and *HMR* (Figure 1D), suggesting that all three members of the SIR complex were equally distributed, as expected. In addition to the aggregate methylation signal (% of reads methylated at each position), we analyzed methylation of single adenines on single reads across *HML* and *HMR* (Figure 1E, Figure 1–figure supplement 2B). Sir3-M.EcoGII methylated most strongly near the *E* and *I* silencers and at the *HMLα1/α2* promoter, with lower, but significant, methylation between these sites (Figure 1E, Figure 1–figure supplement 2B). Analysis of single reads revealed a periodicity of methylation across *HML* and *HMR* (Figure 1E, Figure 1–figure supplement 2B). These small regions of higher methylation corresponded to linker regions between nucleosomes (Figure 1–figure supplement 3). Sir3-M.EcoGII did methylate within nucleosome-occupied regions at *HML* and *HMR* but at a lower frequency (Figure 1–figure supplement 3), consistent with *in vitro* studies that use methylation by M.EcoGII or another non-specific adenine methyltransferase, Hia5, as a measurement of chromatin accessibility (Abdulhay et al., 2020; Altemose et al., 2021; Shipony et al., 2020; Stergachis et al., 2020).

### Nucleosome binding was required for spreading, but not recruitment, of Sir3

Recruitment of the SIR complex to silencers is sequence specific. Rap1, Abf1, and ORC bind at these recruitment sites and recruit the Sir proteins directly through protein-protein interactions. In contrast, SIR complex binding away from recruitment sites is sequence independent and instead relies on interactions with nucleosomes and among the Sir proteins themselves.

We therefore hypothesized that the interaction between Sir3 and nucleosomes would not be required for nucleation at recruitment sites but would be required for binding outside of those recruitment sites. Sir3 has multiple recognized domains (Figure 2A): The BAH domain, which interacts with nucleosomes (Armache et al., 2011; Buchberger et al., 2008; Norris et al., 2008; Onishi et al., 2007; Rudner et al., 2005; Sampath et al., 2009), the AAA^+^ domain, which interacts with Sir4 (Ehrentraut et al., 2011; King et al., 2006; Samel et al., 2017), and the winged helix (wH) domain, which allows for homodimerization (King et al., 2006; Liaw and Lustig, 2006; Oppikofer et al., 2013). We deleted the BAH domain of Sir3-M.EcoGII (*bahΔ*), which largely abrogates Sir3-nucleosome interactions *in vitro* (Buchberger et al., 2008; Onishi et al., 2007; Sampath et al., 2009). Previous studies established that deletion of the BAH domain causes a phenotypic loss of silencing, likely due to a loss of interaction with nucleosomes (Buchberger et al., 2008; Gotta et al., 1998; Onishi et al., 2007), but did not characterize its binding at regions of heterochromatin. Importantly for what follows, deletion of the BAH domain did not destabilize Sir3 (Figure 2B). We also confirmed that *sir3-bahΔ-M.ECOGII* strains displayed a loss of silencing, but found that the loss was more severe at *HML* than at *HMR* (Figure 2C).

**Figure 2.**
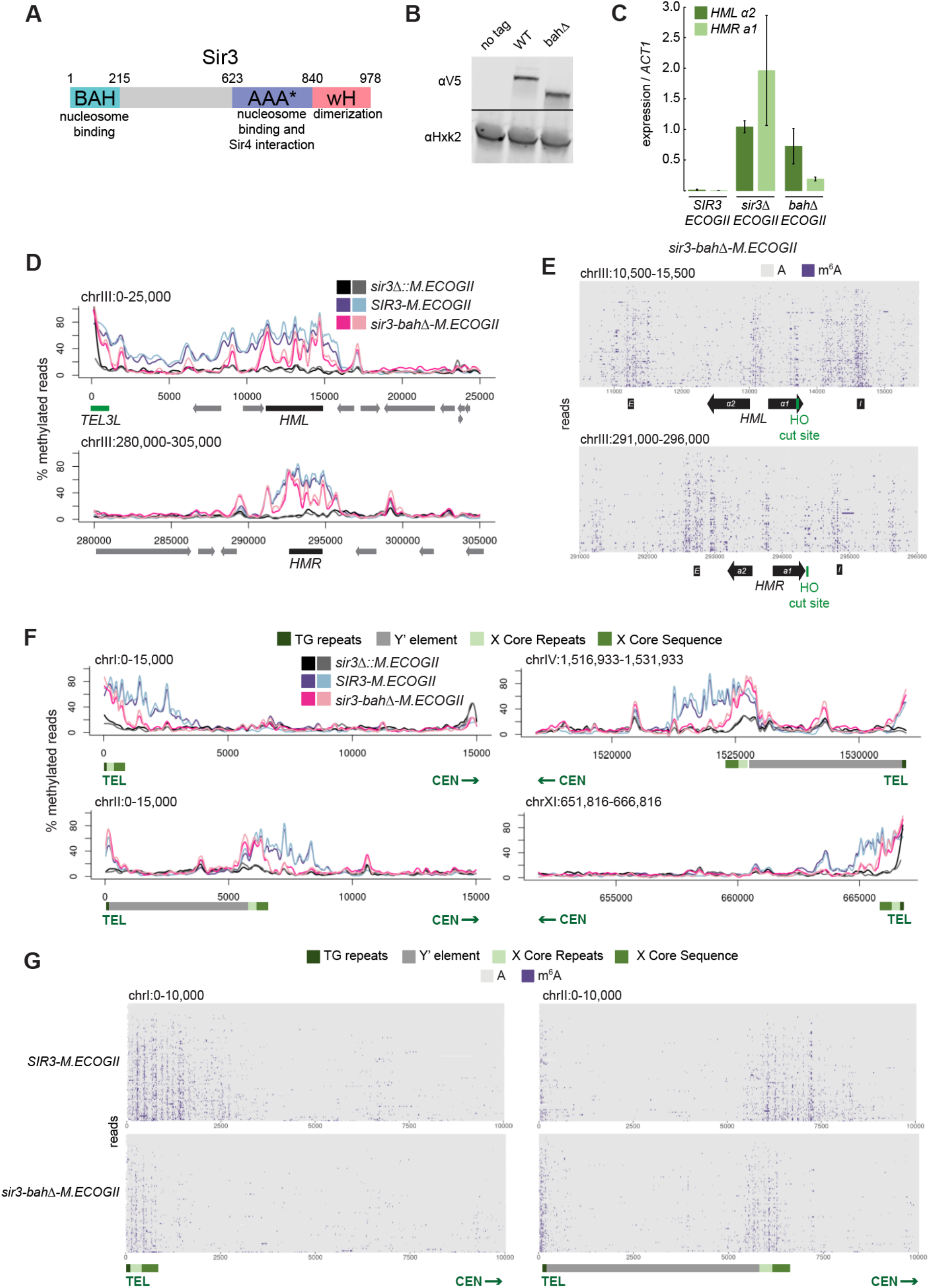
Nucleosome binding was required for spread, but not recruitment, of Sir3 to regions of heterochromatin. **A)** Schematic of Sir3 protein domains. **B)** Protein immunoblotting in strains expressing Sir3 (no tag, JRY11699), Sir3-3xV5 (JRY12601), and sir3-bahΔ-3xV5 (JRY13621). Top row are 3xV5-tagged Sir3 proteins, and bottom row is the loading control Hxk2. The unedited blot is in Figure 2–Source data 1. **C)** RT-qPCR of *HMLα2* and *HMRa1* mRNA, normalized to *ACT1* mRNA, in strains expressing *SIR3-M.ECOGII* (JRY12840, JRY13027), *sir3Δ::M.ECOGII* (JRY13029, JRY13030), and *sir3-bahΔ-M.ECOGII* (JRY13438, JRY13439). Bars are the average of three biological replicates, and bars mark one standard deviation. **D)** Aggregate methylation results at *HML* (top) and *HMR* (bottom) from long-read Nanopore sequencing of *sir3Δ::M.ECOGII* (JRY13029, JRY13030), *SIR3-M.ECOGII* (JRY12840, JRY13027), and *sir3-bahΔ-M.ECOGII* (JRY13438). Plots are as described in Fig 1D. **E)** Single-read plots from long-read nanopore sequencing of *sir3-bahΔ-M.ECOGII* (JRY13438) at *HML* (top) and *HMR* (bottom). Plots are as described in Fig 1E. **F)** Aggregate methylation results at four representative telomeres (1L, 2L, 4R, and 11R) from long-read Nanopore sequencing of the same strains as Fig 2D. Shown are 15 kb windows of each telomere. Plots are as described in Fig 1D. **G)** Single-read plots from long-read nanopore sequencing of *SIR3-M.ECOGII* (JRY13027) and *sir3-bahΔ-M.ECOGII* (JRY13438) at two representative telomeres (1L and 2L). Shown are 10 kb windows of each telomere.

Despite the loss of silencing at *HML* and *HMR* in the *sir3-bahΔ-M.ECOGII* strain, there was still detectable methylation across the two loci (Figure 2D, 2E, Figure 2–figure supplement 1A). At the aggregate level, sir3-bahΔ-M.EcoGII methylated silencers at *HML* and *HMR* at the same level as Sir3-M.EcoGII but showed decreased methylation between them (Figure 2D). Analysis of single reads spanning *HML* and *HMR* in the *bahΔ* mutant showed similarly strong levels of methylation at silencers, and revealed strong methylation both at the promoters of *HML* and *HMR* and the recognition site for the HO endonuclease (Figure 2E). In contrast, little methylation was seen over gene bodies between these sites (Figure 2E).

The technology’s long-read capacity also allowed analysis of the repetitive and highly homologous telomeres. In addition to methylating *HML* and *HMR* (Figure 1), Sir3-M.EcoGII strongly methylated telomeres at TG repeats and X elements (Figure 2F, Figure 2–figure supplement 1B, Figure 2–figure supplement 2), and the periodicity of methylation was apparent on single reads as well (Figure 2G), likely corresponding to more-accessible linker regions. The loss of binding outside of recruitment sites of sir3-bahΔ-M.EcoGII was more striking at telomeres than *HML* and *HMR*, where methylation by the *bahΔ* mutant matched wild-type levels at TG and X repeats but dropped off steeply centromere-proximal to the X elements (Figure 2F, 2G, Figure 2–figure supplement 1B, Figure 2–figure supplement 2). The results at telomeres, supported by the data at *HML* and *HMR*, suggested that the nucleosome binding activity of Sir3 was required for Sir3 to spread away from recruitment sites, but not for its initial recruitment.

### *SIR3* expression level did not limit its spread from recruitment sites

In addition to understanding what enables Sir3 spreading, we were also interested in what limits its spread. One common feature of heterochromatin proteins is that their activity is dose-dependent: lowered expression causes loss of heterochromatin whereas elevated expression can cause silencing of genes near heterochromatin (Henikoff, 1996; Locke et al., 1988). Indeed, it was previously reported that overexpression of Sir3 results in its spread beyond wild-type boundaries, accompanied by repression of genes in those extended regions (Hecht et al., 1996; Hocher et al., 2018; Ng et al., 2003; Renauld et al., 1993; Strahl-Bolsinger et al., 1997). To provide an independent test of those conclusions, we tested whether the expression level of Sir3 limited how far it could spread beyond recruitment sites at *HML*, *HMR*, and telomeres. Expression of both *SIR3-M.ECOGII* from a multi-copy 2μ plasmid was 10-fold higher than the endogenous level of *SIR3-M.ECOGII* in a strain carrying an empty 2μ plasmid (Figure 3A). Overexpression of *SIR3-M.ECOGII* had no effect on silencing at *HML* and *HMR* (Figure 3B).

**Figure 3.**
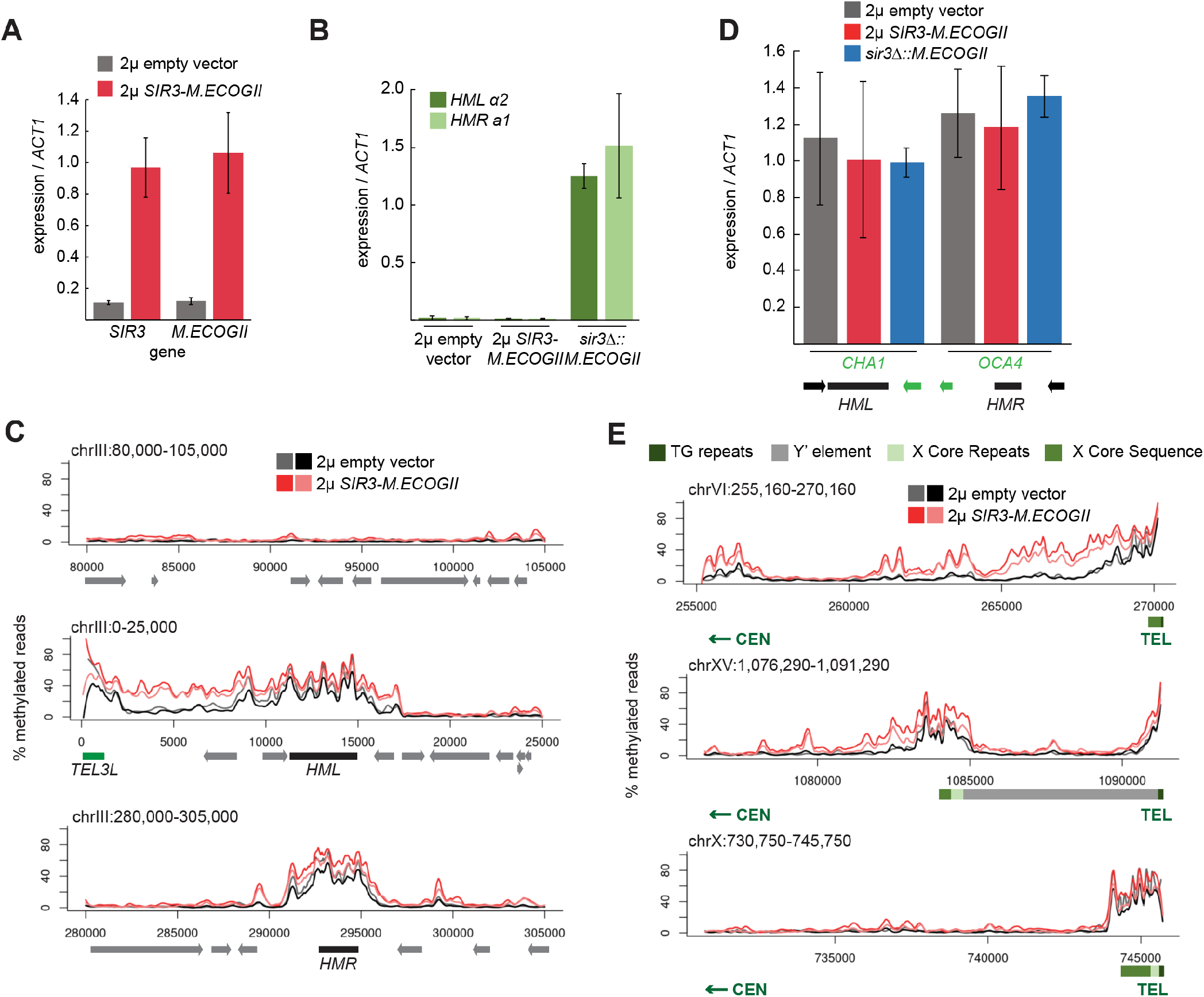
Sir3 expression was not limiting for its spread from recruitment sites. **A)** RT-qPCR of *SIR3* and *M.ECOGII* mRNA normalized to *ACT1* mRNA in strains carrying an empty multi-copy 2μ vector (JRY13670, JRY13671) and strains carrying a multi-copy 2μ *SIR3-M.ECOGII* plasmid (JRY13672, JRY13673). Data are the average of four biological replicates, and bars mark one standard deviation. **B)** RT-qPCR of *HMLα2* and *HMRa1* mRNA normalized to *ACT1* mRNA in the same strains as Fig 3A as well as *sir3Δ::M.ECOGII* (JRY13029, JRY13030). Data are the average of four biological replicates, and bars mark one standard deviation. **C)** Aggregate methylation results at a control region on chromosome III to show background levels of methylation (top row), at *HML* (middle row) and *HMR* (bottom row) from long-read Nanopore sequencing of the same strains in Fig 3A. The two colors for each genotype correspond to two biological replicates. Plots are as described in Fig 1D. **D)** RT-qPCR of CHA1 and OCA4 mRNA normalized to ACT1 mRNA in the same strains as Fig 3B. **E)** Aggregate methylation results at three representative telomeres (6R, 15R, and 10R) from long-read Nanopore sequencing of the same strains as Fig 3A. The two colors for each genotype correspond to two biological replicates Shown are 15 kb windows of each telomere. Plots are as described in Fig 1D.

Overexpression of *SIR3-M.ECOGII* had little effect on the boundaries of methylation at *HML* and *HMR*. Strains overexpressing *SIR3-M.ECOGII* had increased methylation over both loci but no new methylation outside the bounds of strains expressing only one copy of *SIR3-M.ECOGII* (Figure 3C). There was some increase in methylation over the promoters of two genes closest to *HML* and *HMR* (*CHA1* and *OCA4*, respectively), but it did not result in any change in the level of their expression (Figure 3D).

Surprisingly, the results at telomeres were qualitatively similar but revealed three categories of effects. Some telomeres showed a large increase in the amount of methylation upon overexpression of *SIR3-M.ECOGII* with a small extension of binding farther into the chromosome (Figure 3E, top row, Figure 3–figure supplement 1). Some telomeres showed a modest increase in the amount methylation upon overexpression of *SIR3-M.ECOGII* with little, if any, extension of range (Figure 3E, middle row, Figure 3–figure supplement 1). Finally, some telomeres showed no appreciable change in methylation (Figure 3E, bottom row, Figure 3–figure supplement 1). Only three of the 32 telomeres, including the paradigmatic telomere 6R from earlier studies, showed convincing spread of methylation to new sites compared to telomeres in strains expressing one copy of *SIR3-M.ECOGII* (Figure 3E, Figure 3–figure supplement 1). Therefore, the expression level of *SIR3* was not a universal limiting factor in heterochromatin spread. These results imply the existence of other chromatin features that create boundaries for Sir3 spreading.

### Repression of *HML* and *HMR* preceded heterochromatin maturation

To evaluate the dynamics of Sir3 recruitment and spreading during the establishment of heterochromatin over time, we used a temperature-sensitive allele of *SIR3*, *sir3-8,* fused to *M.ECOGII* and took samples at various time points for Nanopore sequencing after switching from the restrictive (37°C) to the permissive (25°C) temperature (Figure 4A). In agreement with previous studies (Stone et al., 2000), growth at 37°C caused lower protein levels of *sir3-8* (Figure 4B). Over the course of 150 minutes, the protein levels of *sir3-8* slowly increased to match levels in constitutive 25°C growth conditions (Figure 4B). In cells grown at 37°C, *sir3-8-M.ECOGII* did not methylate *HML* and *HMR*, but when grown constitutively at 25°C, there was strong methylation over both loci (Figure 4C, Figure 4–figure supplement 1).

**Figure 4.**
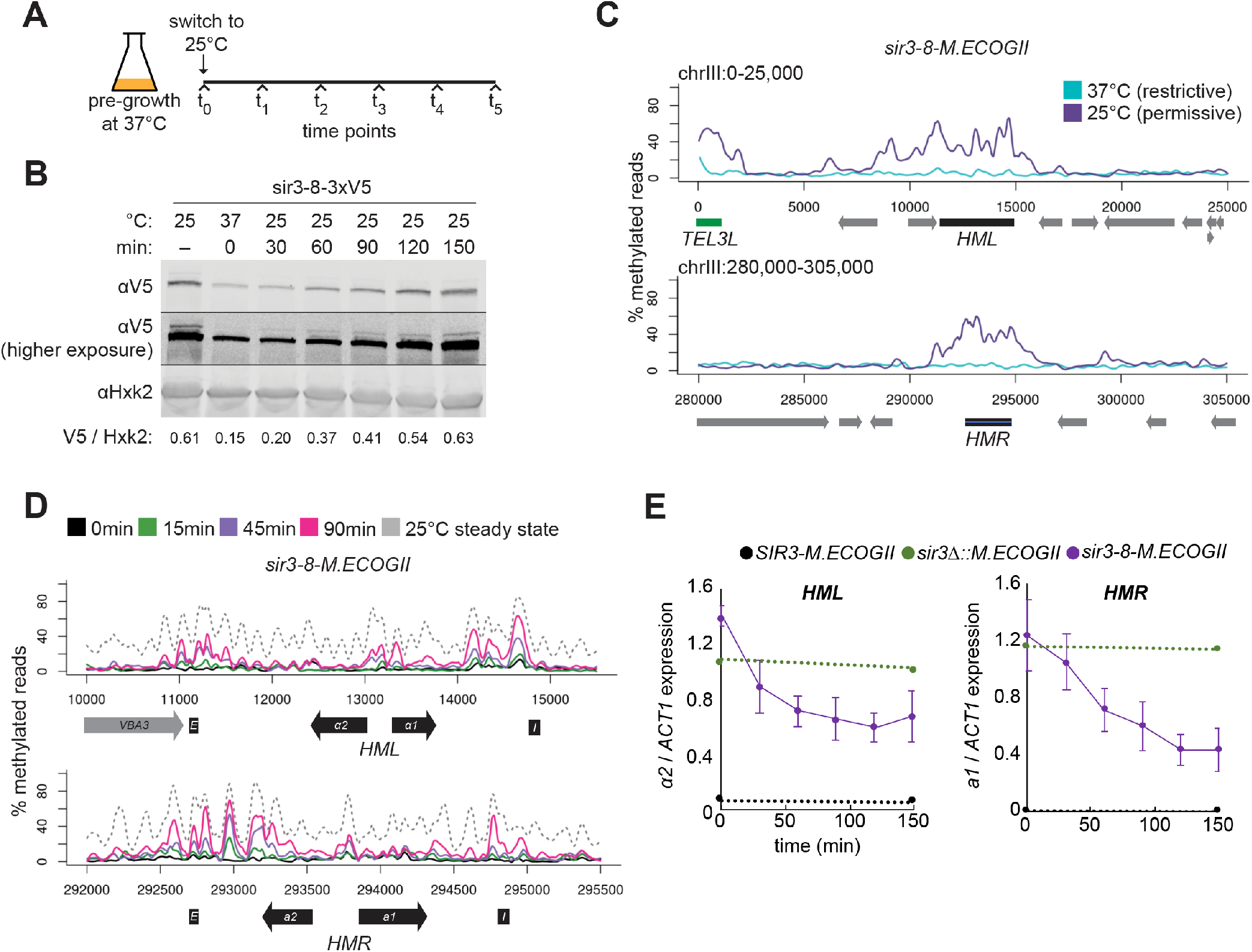
Repression of *HML* and *HMR* preceded heterochromatin maturation. **A)** Schematic of temperature-shift time course with *sir3-8-M.ECOGII*. **B)** Protein immunoblotting in a strain expressing sir3-8-3xV5 (JRY13467) constitutively at 25°C (first lane), constitutively at 37°C (second lane), and at 30 min, 60 min, 90 min, 120 min, and 150 min after a shift to 25°C. Top row is 3xV5-tagged sir3-8 protein, the middle row is the same as the top row but at a higher exposure, and the bottom row is the loading control Hxk2. The unedited blot is in Figure 4–Source data 1. **C)** Aggregate methylation results at *HML* (top) and *HMR* (bottom) from long-read Nanopore sequencing of strains expressing *sir3-8-M.ECOGII* (JRY13114) grown constitutively at 25°C or 37°C. Plots are as described in Fig 1D. **D)** Aggregate methylation results at *HML* (top) and *HMR* (bottom) from long-read Nanopore sequencing of a strain expressing *sir3-8-M.ECOGII* (JRY13134) grown constitutively at 25°C (dotted gray line) and collected at 0 min, 15 min, 45 min, and 90 min after a temperature switch from 37°C to 25°C. **E)** RT-qPCR of *HMLα2* (left) and *HMRa1* (right) mRNA in strains expressing *SIR3-M.ECOGII* (black, JRY13027, JRY12840), *sir3Δ::M.ECOGII* (green, JRY13029, JRY13030) or *sir3-8-M.ECOGII* (purple, JRY13114, JRY13134) collected at 0 min, 30 min, 60 min, 90 min, 120 min, and 150 min after a temperature switch from 37°C to 25°C. Points are the average of three biological replicates and bars mark one standard deviation.

Over a 90-minute time course, methylation increased only over the silencers and promoters of *HML* and *HMR* (Figure 4D, Figure 4–figure supplement 2, solid lines). Methylation at the promoter of *HML* during the time course and in the *sir3-bahΔ* mutant was expected, as it contains a Rap1 binding site, and Rap1 interacts directly with Sir3 and Sir4. However, methylation at the promoter of *HMR* at these early time points and in the *sir3-bahΔ* mutant was a surprise, as there is not a known SIR complex-interacting protein that binds at the promoter. In the absence of Sir3, Sir4 is still bound at silencers (Goodnight and Rine, 2020; Hoppe et al., 2002; Rusche et al., 2002), so the faster recruitment at these sites, and perhaps the promoters as well, might be due to the interaction of Sir3 with Sir4 and Rap1.

By 90 minutes (~1 cell division), methylation at no position reached the level found in cells constitutively grown at 25°C–the level of methylation of mature, stable heterochromatin (Figure 4D, Figure 4–figure supplement 2, dotted line). Strikingly, even by 30 minutes after the temperature shift, when methylation was just rising above background at silencers, partial repression of *HML* and *HMR* was apparent (Figure 4E). This result suggested that binding of Sir3 at silencers and promoters was sufficient for partial repression and preceded its spread over the entirety of both loci.

## DISCUSSION

In this study, we developed a new method to study the process of recruitment and spread of the *S. cerevisiae* heterochromatin protein Sir3 in living cells with a resolution approximating the frequency of single A-T base pairs. We created a fusion protein between a key structural protein of heterochromatin, Sir3, and the bacterial adenine methyltransferase M.EcoGII that retained function and activity of each. We used DNA methylation as a read-out for Sir3 occupancy on chromatin. Long-read Nanopore sequencing allowed us to distinguish directly between methylated and unmethylated adenine and transcended the limitations of earlier studies imposed by repetitive regions common at telomeres.

The methylation by Sir3-M.EcoGII at *HML* and *HMR* was stronger and had a larger footprint than its occupancy as judged by ChIP-seq, suggesting that our method captured transient contacts of Sir3 with chromatin that ChIP-seq could not. This result reinforced the idea that protein-chromatin interactions are dynamic, even for a feature like heterochromatin, commonly thought of as ‘stable’. We also harnessed the power of single base-pair resolution afforded by Nanopore sequencing to distinguish between recruitment and spread of Sir3 by studying a mutant of Sir3 whose distribution and binding profile had not yet been characterized, *sir3-bahΔ*. This mutant cannot bind to nucleosomes and loses transcriptional silencing at *HML* and *HMR* (Buchberger et al., 2008; Gotta et al., 1998; Onishi et al., 2007). The mutant sir3-bahΔ-M.EcoGII was still recruited to silencers at *HML, HMR,* and to telomeres where it methylated local adenines. However, the mutant did not spread beyond those recruitment sites, unlike wild-type Sir3-M.EcoGII. Our findings supported that recruitment and spread were separate processes that involved different interactions between Sir3 and other proteins at silenced loci and telomeres.

The ability to unambiguously map long reads to telomeres allowed us to challenge historic conclusions about Sir3 dosage-driven heterochromatin spreading. Overexpressing Sir3-M.EcoGII increased the methylation signal where there already was methylation at endogenous levels of expression, but the boundaries where methylation dropped off at *HML*, *HMR,* and telomeres mainly remained fixed in the two conditions. This result suggested that overexpression of Sir3 was not sufficient for spreading past most wild-type boundaries. The original idea that overexpression of Sir3 results in its further spread relied on low-resolution RT-PCR at two telomeres, 5R and 6R (Hecht et al., 1996; Renauld et al., 1993; Strahl-Bolsinger et al., 1997). In our genome-wide analysis, telomere 6R was an exception, not the rule, as most telomeres did not show spreading of Sir3 upon its overexpression. We were not able to reproduce the result at telomere 5R (Hecht et al., 1996).

Our data suggested that binding of Sir3 outside of *HML*, *HMR*, X elements, and telomere TG repeats was probabilistic: When Sir3 was overexpressed, its interactions outside of recruitment sites became more frequent, thus increasing the methylation signal produced by Sir3-M.EcoGII. However, most boundaries were left largely unchanged under overexpression conditions, in agreement with more recent Sir3 ChIP-seq results (Radman-Livaja et al., 2011). Perhaps other features, such as transcription, tRNA genes (Donze and Kamakaka, 2001; Simms et al., 2008; Valenzuela et al., 2009), the presence of histone variants like H2A.Z (Babiarz et al., 2006; Giaimo et al., 2019; Meneghini et al., 2003; Venkatasubrahmanyam et al., 2007), or the presence of Sir3-inhibiting chromatin marks like methylation of H3 on lysine 79 (Altaf et al., 2007; Ng et al., 2003, 2002; Oki et al., 2004; Park et al., 2002; Stulemeijer et al., 2011), enforce a boundary that overexpression of Sir3 itself cannot overcome.

Two results suggested that repression of *HML* and *HMR* did not require that Sir3 occupy the entire locus to the level seen in wild-type cells. During the temperature-switch time course, methylation by sir3-8-M.EcoGII appeared first and most strongly at promoters and silencers of *HML* and *HMR*, with little or no detectable methylation between these sites. Yet, partial transcriptional repression was already evident at both loci within 30 minutes. The gradual repression that appeared during the time course was consistent with single-cell studies that show gradual tuning down of transcription during silencing establishment (Goodnight and Rine, 2020). The level of repression achieved during the time course was commensurate with that expected within the first cell cycle following restoration of Sir3 function (Goodnight and Rine, 2020). We also found that *sir3-bahΔ*, a nucleosome-binding mutant that was recruited to silencers and promoters but could not bind outside of them, achieved some repression at both *HML* and *HMR*. Repression was stronger at *HMR* than at *HML* in the *bahΔ* mutant, perhaps because *HMR* is smaller and less dependent on spreading. The time course and *bahΔ* results together suggested a difference between repression of transcription per se and the stability of silencing ultimately achieved by the SIR complex binding over the entirety of *HML* and *HMR.*

We focused our efforts on the heterochromatin protein Sir3 in *S. cerevisiae,* but this method may have broad utility. By relaxing the requirement for stable binding to detect interaction of a protein with DNA or chromatin, the method could find binding sites of transcription factors and/or chromatin-binding proteins that elude detection by ChIP-seq. Further, with an inducible M.EcoGII fusion protein, one could track processes over time like the spread of heterochromatin proteins during X-chromosome inactivation, the movement of cohesin and condensin along chromosomes during chromosome pairing and condensation, and the homology search during homologous recombination. Improvements in Nanopore technology and modified base-calling software will likely extend the method’s utility to other processes that have directional movement along chromosomes.

## FIGURE SUPPLEMENTS

### Figure 1 Supplements

**Figure 1–Figure Supplement 1.**
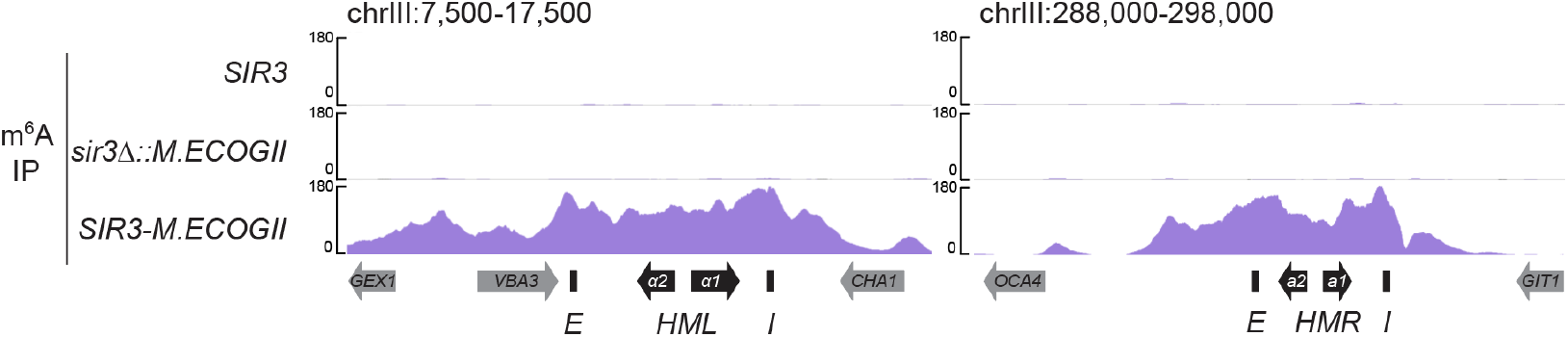
*Sir3-M.EcoGII* strongly and specifically methylated *HML* and *HMR*. Biological replicates of DIP-seq of no EcoGII (top row, JRY09316), *sir3Δ::M.ECOGII* (middle row, JRY12838), and *SIR3-M.ECOGII* (bottom row, JRY13027)

**Figure 1–Figure Supplement 2.**
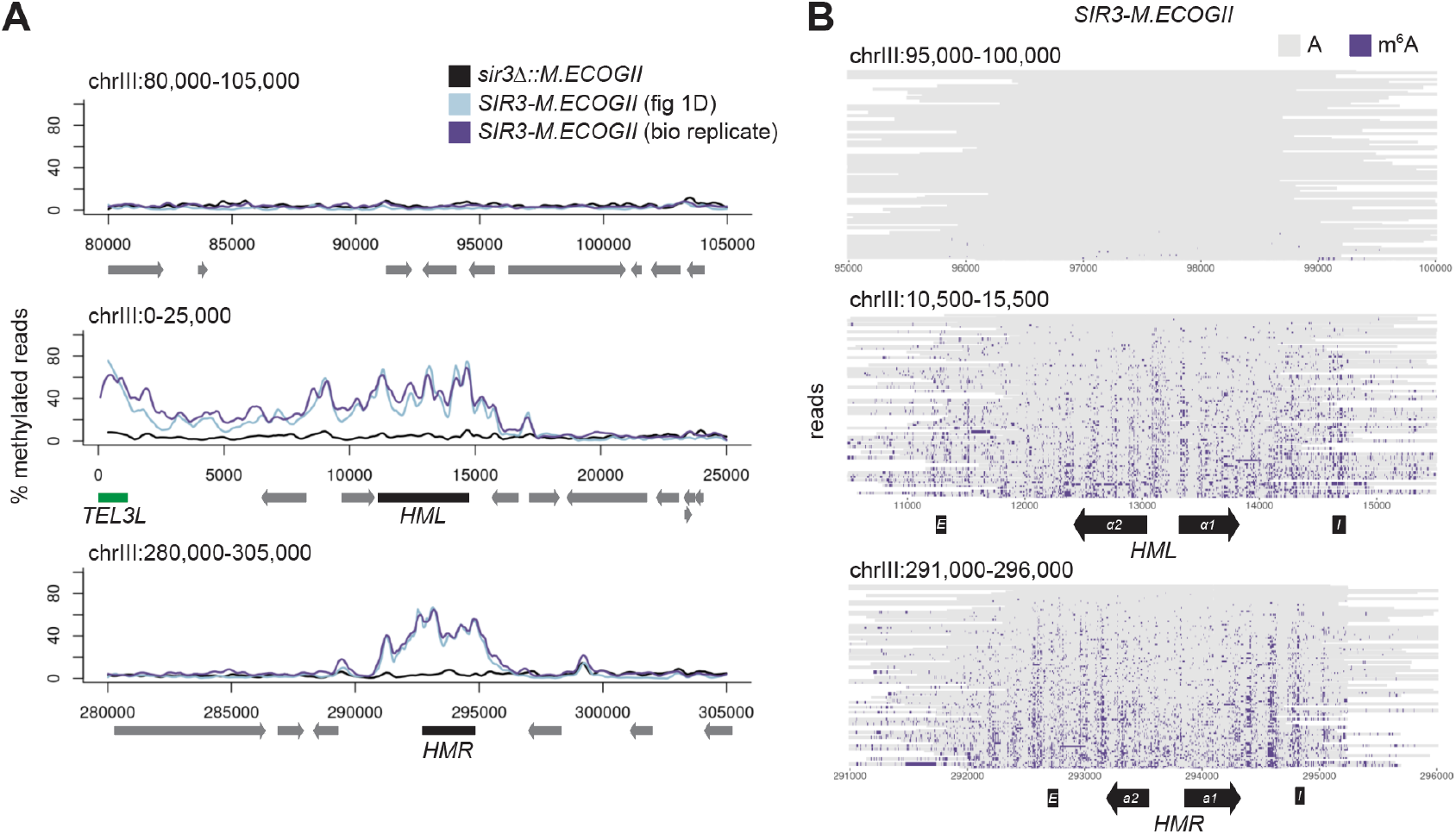
*Sir3-M.EcoGII* strongly and specifically methylated *HML* and *HMR*. **A)** Aggregate results from long-read Nanopore sequencing of *sir3Δ::M.ECOGII* (black line, JRY12838, same as figure 1D), *SIR3-M.ECOGII* (blue line, JRY13027, same as figure 1D), and a biological replicate of *SIR3-M.ECOGII* (purple line, JRY12840). Plots are as described in Fig 1D. **B)** Single-read plots from a biological replicate of long-read Nanopore sequencing of *SIR3-M.ECOGII* (JRY13027). Plots are as described in Fig 1E.

**Figure 1–Figure Supplement 3.**
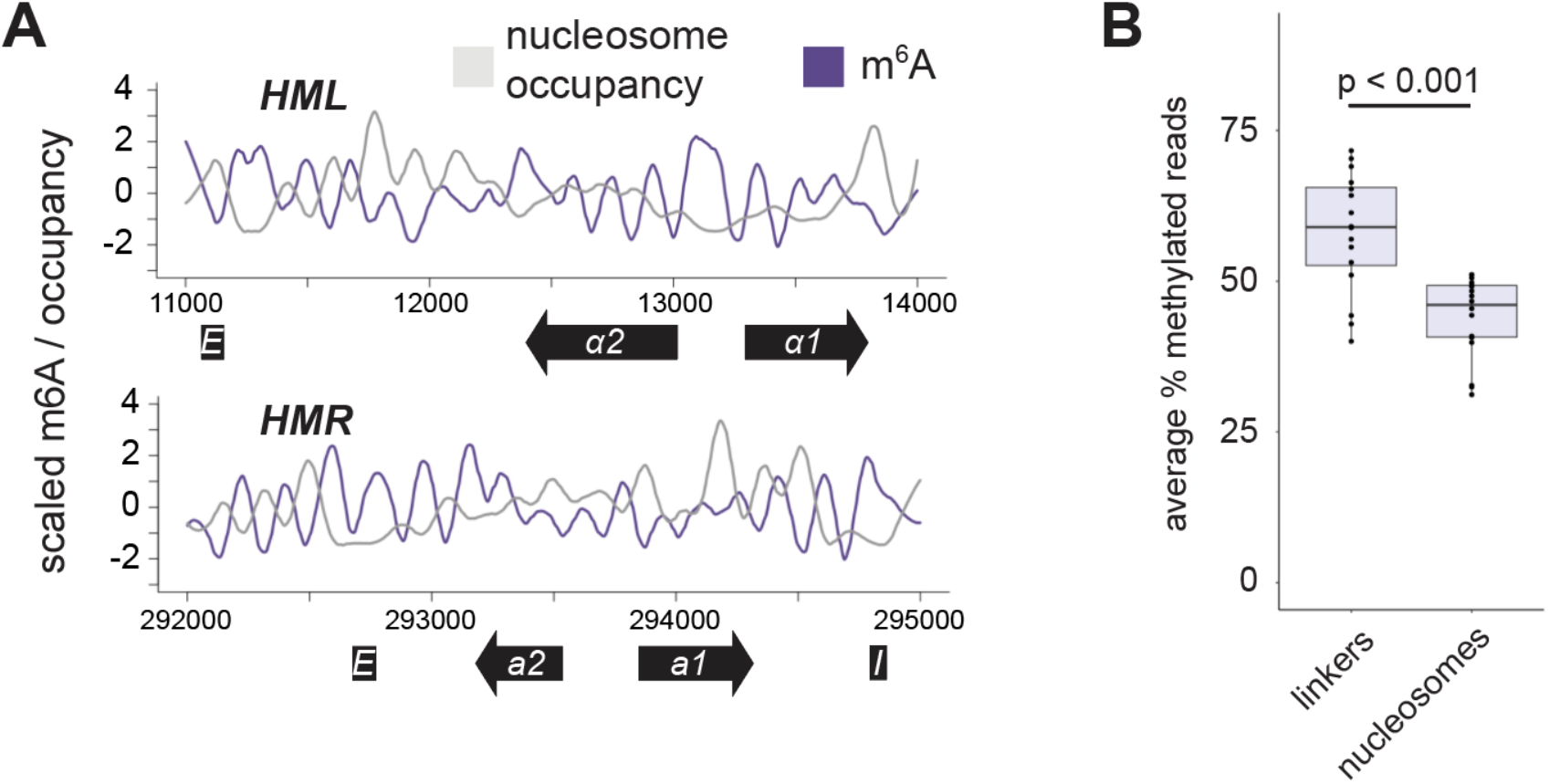
Sir3-M.EcoGII preferentially methylated linker regions. **A)**Aggregate methylation signal by Nanopore sequencing (purple line) of *SIR3-M.ECOGII* (JRY13027) plotted against nucleosome occupancy (gray line, Chereji et al. 2018) over a 3 kb window at *HML* (top) and *HMR* (bottom). The y-axis represents the signal of both % of reads methylated and nucleosome occupancy scaled to fit in the same plot. Data were smoothed using a loess function. **B)** The average % of methylated reads from *SIR3-M.ECOGII* (JRY13027) in linker regions at *HML* and *HMR* (nucleosome occupancy below a threshold of 0.4 from Chereji et al. 2018) compared to nucleosomal regions at *HML* and *HMR* (nucleosome occupancy above a threshold of 0.4 from Chereji et al. 2018). The center line of each box plot represents the median. The boxes represent the 25th and 75th percentiles. Whiskers represent the range of values within 1.5× the interquartile range. P-value was calculated using a quasibinomial general linear model (glm, see Source Code 5).

### Figure 2 Supplements

**Figure 2–Figure Supplement 1.**
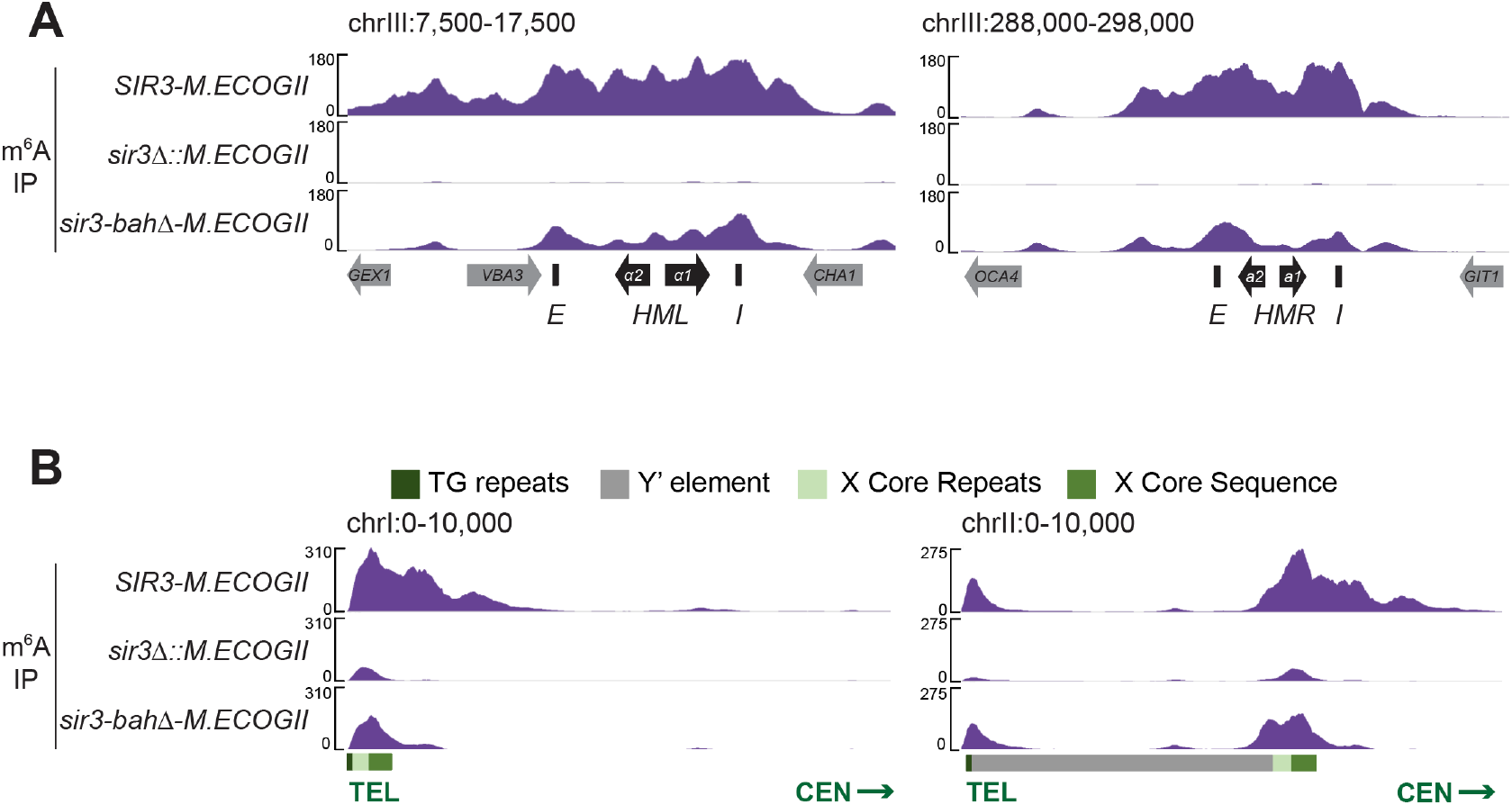
DIP-seq of *SIR3-M.ECOGII* (top row, JRY13027), *sir3Δ::M.ECOGII* (middle row, JRY13030), and *sir3-bahΔ-M.ECOGII* (bottom row, JRY13438) Input results are plotted but not visible due to the strong DIP-seq signals. **A)** Shown are 10 kb regions centered at *HML* (left) and *HMR* (right). **B)** Shown are 10 kb regions at two representative telomeres (1L and 2L)

**Figure 2–Figure Supplement 2.**
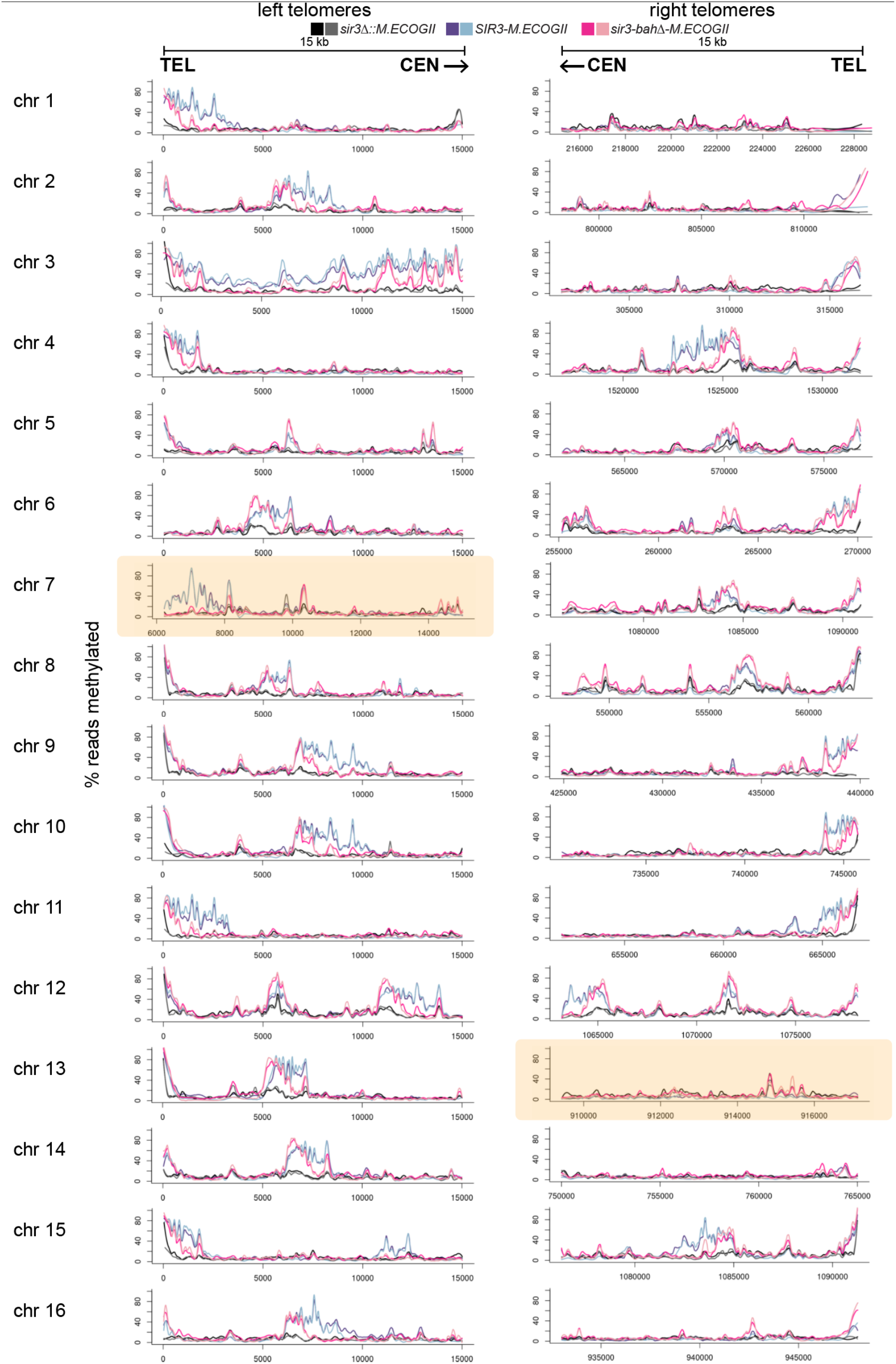
Methylation by Sir3-M.EcoGII and sir3-bahΔ-M.EcoGII at all 32 telomeres. Aggregate methylation results from long-read Nanopore sequencing of the same strains as Fig 2D. Shown are 15 kb windows of each telomere. Plots are as described in Fig 1D. Highlighted in yellow are windows shorter than 15kb due to discrepancies between the S288C and W303 genomes (see Ellahi et al. 2015)

### Figure 3 Supplement

**Figure 3–Figure Supplement 1.**
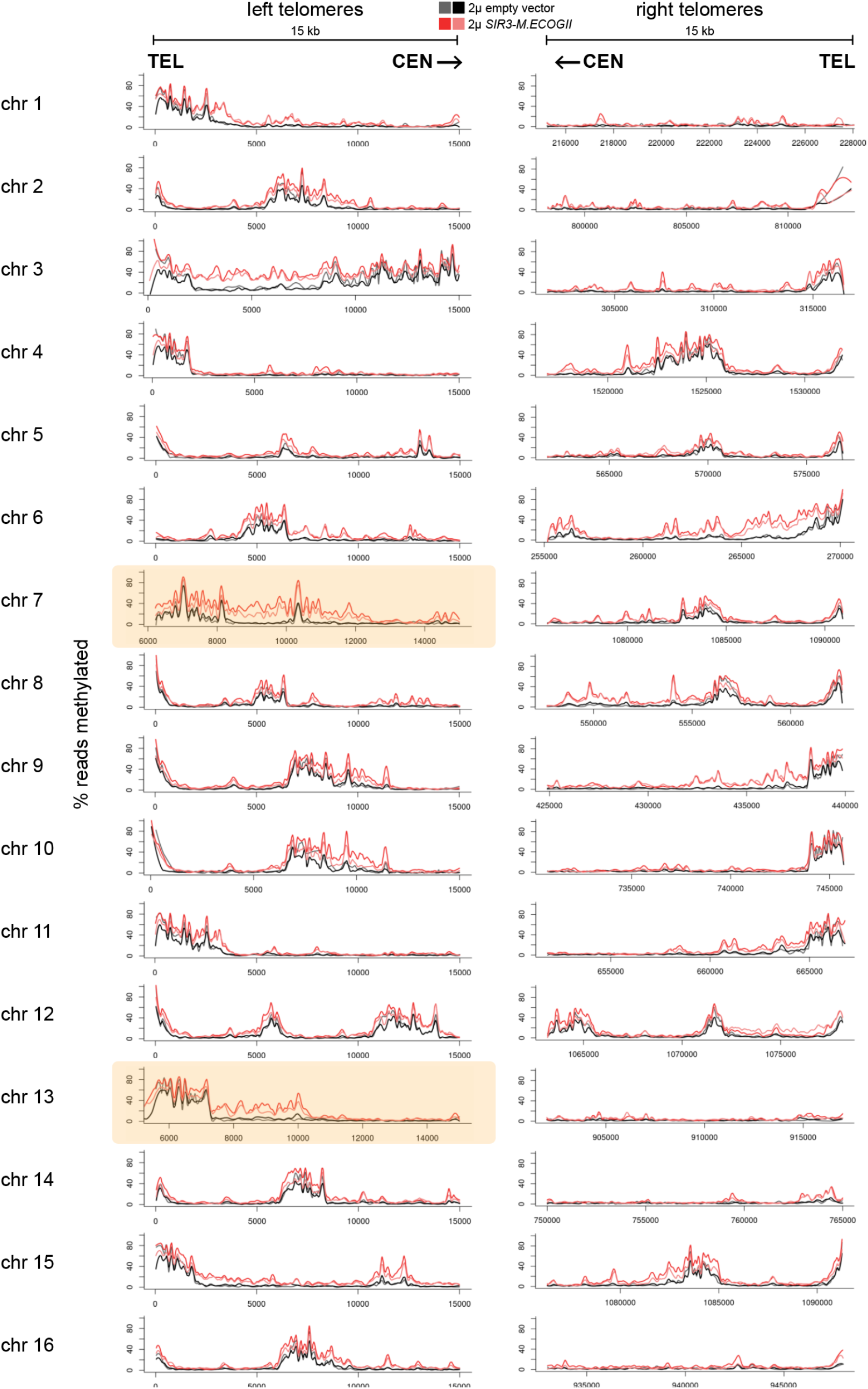
Methylation upon overexpression of *SIR3-M.ECOGII* at all 32 telomeres. Aggregate methylation results from long-read Nanopore sequencing of the same strains as Fig 3B. Shown are 15 kb windows of each telomere. Plots are as described in Fig 1D. Highlighted in yellow are windows shorter than 15kb due to discrepancies between the S288C and W303 genomes (see Ellahi et al. 2015)

### Figure 4 Supplements

**Figure 4–Figure Supplement 1.**
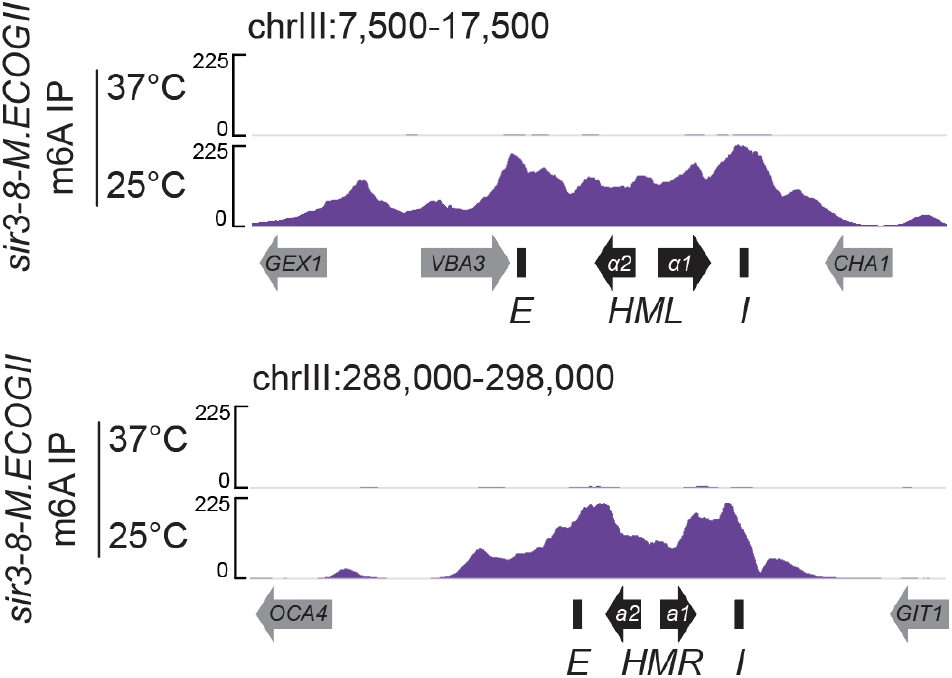
DIP-seq of *sir3-8-M.ECOGII* (JRY13114) Shown are 10 kb regions centered at *HML* (left) and *HMR* (right). Cells were grown constitutively at either 25°C or 37°C. Input results are plotted but not visible due to the strong DIP-seq signals.

**Figure 4–Figure Supplement 2.**
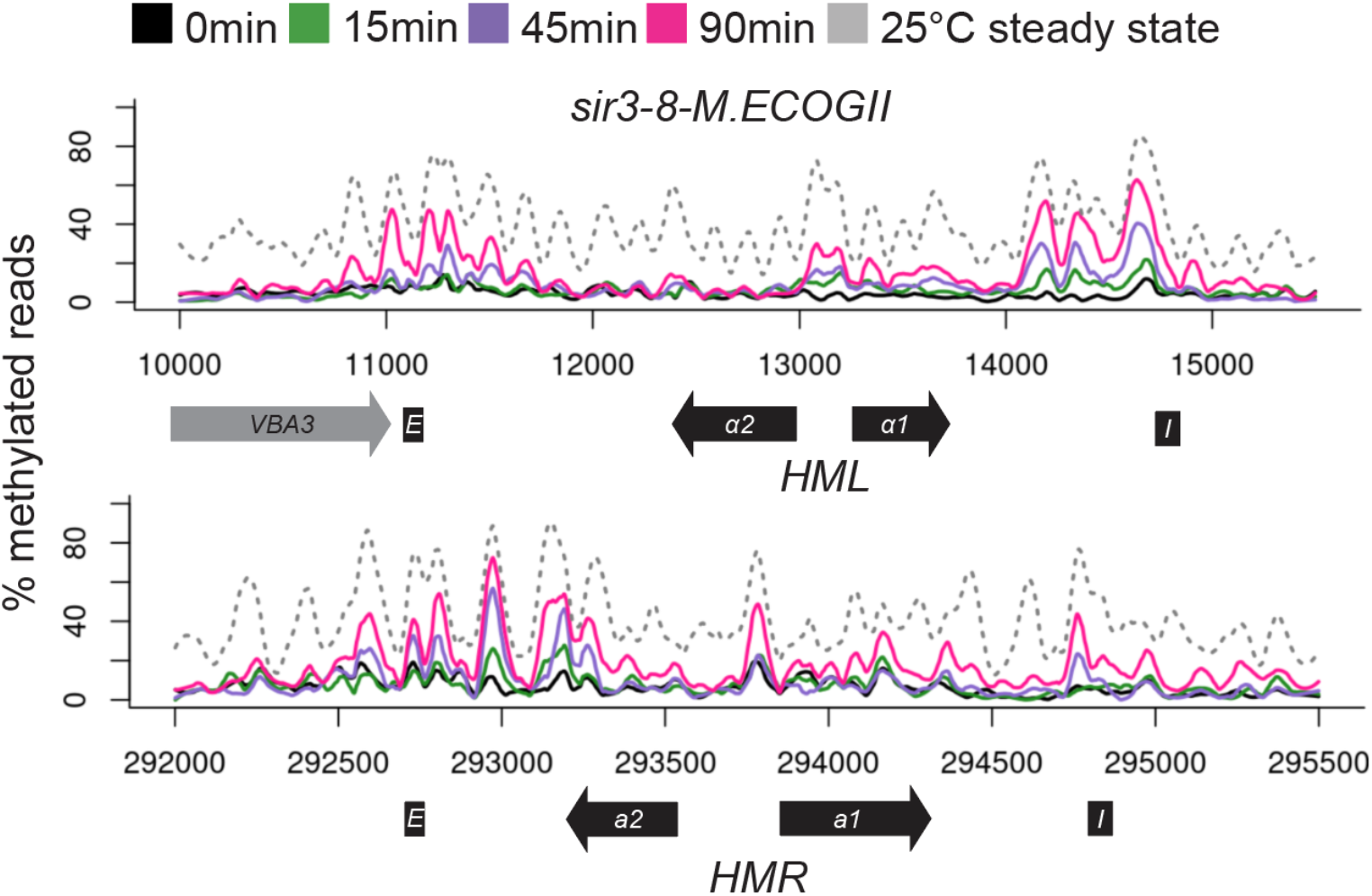
Nanopore sequencing over temperature switch time course (biological replicate) Aggregate methylation results at *HML* (top) and *HMR* (bottom) from long-read Nanopore sequencing of a strain expressing *sir3-8-M.ECOGII* (JRY13114) grown constitutively at 25°C (dotted gray line) and collected at 0 min, 15 min, 45 min, and 90 min after a temperature switch from 37°C to 25°C.

## MATERIALS AND METHODS

### Strains

All strains in this study were derived from W303 (Supplementary File 1). For biological replicates, independent cultures were started from the same strain or isogenic strains as indicated by strain numbers in figure legends. *M.ECOGII* integrations were created by one-step integration of a PCR-amplified *M.ECOGII-natMX* cassette from pJR3525 using the primers listed in Supplementary File 3. Deletion of the BAH domain of *SIR3* was done using CRISPR-Cas9 gene editing using the guide RNA and repair template listed in Supplementary File 3. The guide RNA target and nontarget strands were integrated into a single guide RNA dropout-Cas9 expression plasmid (pJR3428, (Brothers and Rine, 2019) by Golden Gate cloning, using the restriction enzyme *Bsm*BI as described in (Lee et al., 2015). The repair template was made by annealing oligos described in Supplementary File 3 and extending the 3’ ends using Phusion Polymerase (New England Biolabs, Beverly, MA). *SIR3* overexpression strain and its control strain was created by transformation and maintenance of 2-micron plasmids pJR3526 and YEp24, respectively.

### Plasmids

Plasmids used in this study are listed in Supplementary File 2. The *M.ECOGII-NatMX6* tagging plasmid (pJR3525) was made using standard Gibson cloning into pFA6a-natMX6 (Hentges et al., 2005). The codon-optimized *M.ECOGII* ORF with homology to the vector backbone was on a gene block (Supplementary File 3) from Integrated DNA Technologies (IDT, Coralville, IA) and was inserted into pFA6a-natMX6 linearized by PCR. The *SIR3p-SIR3-M.ECOGII* overexpression plasmid (pJR3526) was made using standard Gibson cloning into YEp24, a 2μ yeast expression plasmid carrying a *URA3* selectable marker. The sequences for each plasmid are available in Supplementary Materials as fasta and .dna files.

### Growth and fluorescence imaging of colonies

Strains were grown in a patch on YPD overnight at 30°C. Cells were then resuspended in water and plated for single colonies on a YPD plate. Colonies were imaged after 3 days of growth at 30°C. At least 10 colonies per genotype were imaged using a Leica M205FA fluorescence stereomicroscope, a Leica DFC3000G CCD camera, and a Plan Apo x0.63 objective. All colonies were imaged at a magnification of 10X. Image analysis and assembly was performed using Fiji software (Schindelin et al., 2012).

### ChIP-seq

#### Sample Collection

Strains were grown to mid-log phase in YPD at 30°C. Approximately 10^9^ cells were collected, washed, and fixed for 15 minutes at 30°C in a final concentration of 1% formaldehyde. Fixation was quenched with a final concentration of 300mM glycine for 10 minutes at 30°C. Cells were washed 1X with PBS and 2X with FA lysis buffer (50mM HEPES pH7.5, 150 mM NaCl, 1 mM EDTA, 1% v/v Triton X-100, 0.1% w/v sodium deoxycholate, 0.1% SDS) before flash freezing pellets in a 2 mL screw-cap tube. Pellets were resuspended in 1 mL FA lysis buffer and 500 μL of 0.5 mm zirconium ceramic beads (BioSpec Products, Bartlesville, OK, Cat # 11079105z) were added. Resuspended cells were bead beat with in a FastPrep-24 (MP Biomedicals, Burlingame, CA) at 5.5 amplitude, 4 cycles of 40 sec ON/2 min on ice. Each tube was punctured at the bottom with a hot 20G needle and placed into a new 1.5 mL tube, and sample was spun out of the tube into the new tube by spinning at 150 xg for 1 min. The sample was moved into a 15 mL Bioruptor sonication conical tube with 100 μL of Bioruptor sonication beads (Diagenode, Denville, NJ, Cat # C01020031) and sonicated using the Bioruptor Pico (Diagenode) for 10 cycles of 30 sec ON / 30 sec OFF.

#### Immunoprecipitation

The sonicated extract was moved into a new 1.5 mL tube and spun at 16,000 xg for 15 min at 4°C. The supernatant was moved into a new 1.5mL tube and adjusted to 1 mL volume with FA lysis buffer. 50 μL of sample was taken aside as input, and then 25 μL of 20 mg/mL BSA and 5 μL of anti-V5 antibody (Invitrogen, Waltham, MA, Cat # R960-25) was added to the rest of the sample and rotated overnight at 4°C. 50 μL of Protein A magnetic Dynabeads (Thermo-Fisher, Waltham, MA, Cat # 10002D) were added to the sample and rotated at 4°C for 1 hr. Magnetic beads were immobilized using a magnetic rack and washed by resuspension in 1mL of various buffers in the following order: FA lysis buffer + 0.05% Tween-20, FA lysis buffer + 0.05% Tween-20 + 0.25 mM NaCl, ChIP wash buffer (10 mM Tris pH 8.0, 0.25 M LiCl, 1mM EDTA, 0.5% Nonidet P-40, 0.5% sodium deoxycholate, 0.05% Tween-20), and TE (10 mM Tris pH 8.0, 1 mM EDTA) + 0.05% Tween-20. The washed beads were resuspended in 130 μL of ChIP elution buffer (10 mM Tris pH 7.5, 1 mM EDTA, 1% SDS) and incubated at 65°C shaking at 900rpm overnight. The next day, 2.5 μL of 10 mg/mL Proteinase K (New England Biolabs, Ipswich, MA, Cat # P8107S) and 2.5 μL of 10 mg/mL RNase A (Thermo-Fisher, Cat # EN0531) were added to the elution and incubated at 42°C for 2 hr. Beads were immobilized on a magnetic rack and the supernatant containing the desired DNA to be sequenced was taken and purified using 1X v/v SPRI Select magnetic beads (Beckman Coulter, Brea, CA, Cat # B23317) according to the manufacturer’s instructions.

#### Library Preparation and Sequencing

Samples were prepared for sequencing using NEBNext Ultra II DNA Library Prep Kit for Illumina (New England Biolabs, Cat # E7645) according to the manufacturer’s instructions. Samples were multiplexed using NEBNext Multiplex Oligos for Illumina (New England Biolabs, Cat # E7335/E7500). Library-prepped samples were sequenced on a MiniSeq System (Illumina, San Diego, CA)

#### Analysis

Sequencing reads were aligned to the S288C sacCer3 reference genome (release R64-2-1_20150113, yeastgenome.org), modified to include *matΔ* using Bowtie2 with the options “--local --soft-clipped-unmapped-tlen --no-unal --no-mixed --no-discordant” (Langmead and Salzberg, 2012). Reads were normalized to the genome-wide median, excluding rDNA, chromosome III, and subtelomeric regions (the first and last 10 kb of each chromosome). Analysis was performed using custom Python scripts modified from (Goodnight and Rine, 2020) (Source Code 1, Source Code 2) and displayed using IGV (Thorvaldsdóttir et al., 2013).

### DIP-seq

#### DNA extraction

Cells were grown to mid-log phase in YPD, Complete Supplement Mixture (CSM) or CSM without Uracil (Sunrise Science Products, Knoxville, TN) at 30°C. Approximately 10^9^ cells were pelleted by centrifugation at 3200 xg for 2 min, washed with 1 mL of water, moved to a 2 mL screw-cap tube, and flash frozen. Cells were resuspended in 400 μL of Triton SDS Lysis Buffer (10 mM Tris pH 8.0, 100 mM NaCl, 1 mM EDTA, 2% Triton X-100, 1% SDS), and 400 μL of phenol:chloroform:isoamyl alcohol 25:24:1 and 300 μL of 0.5 mm zirconium ceramic beads (BioSpec Products, Cat # 11079105z) were added to the resuspension. Cells were lysed by bead beating with in a FastPrep-24 (MP Biomedicals) at 5.5 amplitude, 4 cycles of 40 sec ON/2 min on ice. The aqueous and organic phases were separated by centrifugation at 21,000 xg for 5 min, and the aqueous phase was moved to a new 1.5 mL tube. 400 μL of chloroform was added, vortexed at top speed for ~ 10 sec, and spun down at 21,000 xg for 5 min to separate the aqueous and organic phases. The aqueous phase was moved to a new 1.5 mL tube, and 1 mL of 100% ethanol was added to precipitate nucleic acids. The sample was incubated at 4°C for 10-15 min and then spun down at 21,000 xg for 2 min to pellet the precipitated nucleic acids. Supernatant was discarded, the pellet was air-dried, and then the pellet was resuspended in 400 μL of TE (10mM Tris-HCl pH 8.0, 1 mM EDTA) + 4 μL of 10 mg/mL RNase A (Thermo-Fisher, Cat # EN0531) and incubated at 37°C for 1 hour. 1 mL of 100% ethanol + 10 μL of 4M ammonium acetate was added to the RNase solution and incubated at 4°C for 10-15 min to precipitate DNA. The precipitate was pelleted by centrifugation at 21,000 xg for 2 min, washed 1X with 70% ethanol, air-dried, and resuspended in 150-300 μL of water.

#### Sonication

DNA concentration was measured using Qubit dsDNA HS reagents (Invitrogen, Cat #Q32854), and 6 μg of DNA was diluted to 20 ng/uL in 300 μL of water in 1.5 mL Bioruptor Pico Microtubes for sonication (Diagenode, Cat # C30010016). DNA was sonicated using a Bioruptor Pico (Diagenode) for 18 cycles of 15 sec ON/90 sec OFF. The sonicated DNA was moved to a new 1.5 mL tube.

#### m^6^A IP

DNA was denatured by incubating at 95°C for 10 min and then immediately placed on ice for 5 min. 200 μL of cold water and 500 μL of cold 5X DIP buffer (50 mM NaPO_4_ pH 7.0, 700 mM NaCl, 0.25% Triton X-100) were added to bring the volume up to 1 mL. 50 μL were taken aside as input. 25 μL of 20 mg/mL BSA and 1.8 μg of antibody (Synaptic Systems rabbit anti-m^6^A, Cat 202-003) were added to the rest of the sample and rotated overnight at 4°C. 50 μL of Protein A magnetic Dynabeads (Thermo-Fisher, Cat # 10002D) were added to the sample and rotated at 4°C for 1 hr. Magnetic beads were immobilized using a magnetic rack and washed by resuspension and rotation for 5 min at 4°C in 1mL of various cold buffers in the following order: 2X with 1X DIP buffer + 0.05% Tween-20, 1X with 1X DIP buffer. For elution, beads were resuspended in 190 μL of DIP Digestion Buffer (50 mM Tris-HCl pH 8.0, 10 mM EDTA, 0.5% SDS) + 10 μL of 10 mg/mL Proteinase K (New England Biolabs, Cat # P8107S). DIP digestion buffer was added to input samples up to 200 μL. Both the input and IP samples were incubated at 50°C for 2 hr and then cleaned up using the Qiaquick PCR Purification Kit (Qiagen, Hilden, Germany, Cat # 28104) and eluted in 35 μL of water.

#### Library Preparation and Sequencing

Samples were prepared for sequencing using Accel-NGS 1S Plus DNA Library Kit (Swift Biosciences, Ann Arbor, MI, Cat # 10024) according to the manufacturer’s instructions. Samples were multiplexed using Swift Single Indexing Primers Set A (Swift Biosciences, Cat # X6024). Library prepped samples were sequenced on a MiniSeq System (Illumina).

#### Analysis

Analysis was done as described in the section on ChIP-seq above.

### Nanopore sequencing

#### DNA extraction

Cells were grown to mid-log phase in YPD, CSM (Sunrise Science Products), or CSM without Uracil at 30°C. Approximately 10^8^ cells were pelleted, washed with 1 mL of water, and pellets were flash frozen. gDNA was extracted using the YeaStar Genomic DNA Kit (Genesee Scientific, San Diego, CA, Cat #11-323) according to the manufacturer’s “Protocol 1”. Specifically, thawed cell pellets were resuspended in 240 μL of YD Digestion Buffer + 10 μL R-Zymolyase and incubated at 30°C for 1 hr. 240 μL of YD Lysis buffer was added to the solution and vortexed at top speed for 15 sec. 500 μL of chloroform was added to the solution and vortexed at top speed for 10 sec and then inverted 10 times. The aqueous and organic phases were separated by centrifugation at 10,000 xg for 2 min, and the aqueous phase was equally separated into two ZymoSpin columns. ZymoSpin columns were spun at 10,000 xg, washed 2X with 300 μL of DNA Wash Buffer, and DNA was eluted from each column with 75 μL of water. Eluates were combined. DNA was sheared to ~15-20kb by spinning through a Covaris g-TUBE (Covaris Inc., Woburn, MA, Cat #520079) at 4200 rpm for 1 min, and repeating 1X with the tube flipped the other way in an Eppendorf Centrifuge 5424, according to the Covaris protocol. DNA was purified and concentrated using 1X v/v SPRI Select beads (Beckman Coulter, Cat # B23317) and eluted in 50 μL of water according to the manufacturer’s instructions. DNA concentration was measured using Qubit dsDNA HS reagents (Invitrogen, Cat # Q32854).

#### Library Preparation and Sequencing

Approximately 1-3 μg of purified, sheared genomic DNA was library prepped using the following reagents: NEB Oxford Nanopore Companion (New England Biolabs, Cat # E7180S), NEB Blunt/TA Ligase Master Mix (New England Biolabs, Cat # M0367), NEBNext Quick Ligation Reaction Master Mix (New England Biolabs, Cat # B6058), Oxford Ligation Sequencing Kit (Oxford Nanopore Technologies, Oxford, United Kingdom, Cat # SQK-LSK109), and the Oxford Native Barcoding Expansion 1-12 (Oxford Nanopore Technologies, Cat # EXP-NBD104). The library was prepared and sequenced according to Oxford Nanopore’s protocol for Ligation Sequencing Kit + Native Barcoding Expansion 1-12. Sequencing was done on a MinION sequencer with v9.4 flow cells (Oxford Nanopore Technolgies, Cat # FLO-MIN106).

#### Analysis

Basecalling was first done using Guppy v5.0.11 using the high-accuracy model (dna_r9.4.1_450bps_hac.cfg), and reads were demultiplexed using guppy_barcoder. Read IDs corresponding to each barcode were extracted and written to a .txt file using a custom Python script (Source Code 3). Reads corresponding to each barcode were aligned to the S288C reference genome (release R64-2-1_20150113, yeastgenome.org, modified to include *matΔ*) and modifications called with Megalodon (https://github.com/nanoporetech/megalodon, v2.3.3) using the all-context rerio model (https://github.com/nanoporetech/rerio, res_dna_r941_min_modbases-all-context_v001.cfg) and the flags --mod-motif “Y A 0”, --files_out “basecalls mod_mappings per_read_mods” and --read-ids-filename “barcodeXX_readIDs.txt” (the file that contained the extracted list of readIDs for a given barcode).

Results were aggregated into .bed files using “megalodon_extras aggregate run”, and these files were used for aggregate nanopore plots. Before plotting, aggregated data was filtered to include only adenines with at least 10X coverage, and lines were smoothed using base R loess() function with enp.target = 100 and weighted by the coverage at each position. Assessment of linker-region preference of Sir3-M.EcoGII used nucleosome-occupancy data from GEO Accession GSE97290 (Chereji et al., 2018).

The per-read database from Megalodon was converted into a .txt file using “megalodon_extras per_read_text modified_bases”. For ease of use in RStudio, the data for each chromosome was extracted into its own .txt file using custom bash and awk scripts (Source Code 4) and these files were used for single-read nanopore plots. The probabilities output by Megalodon were made binary by calling adenines with a >0.8 probability of being methylated as “m6A” and all others “A”.

The R scripts (as .html files) used to create each figure can be found in Source Code 5.

#### Limitations

This method showed possible limitations for some contexts: 1) The expression level of the fusion protein could increase the levels of background methylation. We found this to be true with the Sir2-M.EcoGII fusion protein, which is likely expressed at a higher level than Sir3-M.EcoGII due to the higher level of endogenous Sir2 expression. The level of methylation by Sir2-M.EcoGII in heterochromatin regions was higher than by Sir3-M.EcoGII, and background levels of methylation outside of heterochromatin regions was also higher than by Sir3-M.EcoGII. Importantly, the signal at heterochromatin was evident above even this raised background methylation. 2) Methylation at the level of single reads was variable and spotty, possibly due to at least two contributors. There may be occupancies that are too transient to allow methylation. Secondly, computational limitations for calling modified adenines without a guiding sequence motif meant that lower-confidence (probably < 0.8) m^6^A calls were not considered methylated. At present, qualitative conclusions based on single-read data can be made with confidence, but as nanopore technology improves, single-read data will become more amenable to statistical and spatial analysis. 3) Because this method can capture transient interactions better than methods like ChIP-seq it may overestimate degrees of occupancy unless combined with DIP-seq or ChIP-seq.

### RT-qPCR

#### RNA extraction

Cells were grown to mid-log phase in YPD, CSM (Sunrise Science Products), or CSM–Uracil at 30°C, and RNA was extracted using the Qiagen RNeasy kit (Qiagen, Cat # 74104) according to the manufacturer’s instructions for Purification of Total RNA from Yeast. Briefly, ~6 x 10^7^ cells were resuspended in 600 μL of buffer RLT, 500 μL of 0.5 mm zirconium ceramic beads (BioSpec Products, Cat # 11079105z) were added, and cells were lysed by bead beating with in a FastPrep-24 (MP Biomedicals) at 5.5 amplitude, 3 cycles of 40 sec ON/2 min on ice. Cells were pelleted by spinning at 21,000 xg for 2 min, and the supernatant was moved to a new tube. One volume of 70% ethanol was added to the supernatant and the sample was spun through an RNeasy spin column. The column was washed with 350 μL of buffer RW1, then 10 μL of DNase + 70 μL of buffer RDD (Qiagen, Cat # 79256) were added to the column and incubated for 15 min at room temperature. 500 μL of buffer RW1 was added and spun through the column. The column was then washed with 500 μL of buffer RPE 2X, and RNA was eluted with 80-150 μL of RNase-free water.

#### RT-qPCR

Complementary DNA was synthesized using the SuperScript III First-Strand Synthesis System (Invitrogen, Cat # 18080051) and oligo(dT) primers according to the manufacturer’s protocols. Quantitative PCR of complementary DNA was performed using the DyNAmo HS SYBR Green kit (Thermo-Fisher, Cat # F410L) on an Mx3000P machine (Stratagene, La Jolla, CA) using the primers listed in Supplementary File 3. Standard curves were generated using a 10-fold dilution series of one of the prepared samples.

### Protein Immunoblotting

Each strain was grown to saturation overnight in 5mL YPD. Overnight cultures were diluted to ~2 x 10^5^ cells/mL in fresh YPD, grown to mid-log phase, and ~10^8^ cells were harvested and pelleted. Pellets were resuspended in 1 mL of 5% trichloroacetic acid and incubated at 4°C for 10-30 min. The precipitates were pelleted, washed once with 1mL of 100% acetone, and air-dried. Dried pellets were resuspended in 100 μL of protein breakage buffer (50mM Tris-HCl, pH 7.5, 1mM EDTA, 3mM DTT) and an equal volume of 0.5 mm zirconium ceramic beads (BioSpec Products, Cat # 11079105z) followed by four cycles of 40 sec bead beating / 2 min on ice in a FastPrep-24 (MP Biomedicals). 100 μL of 2X Laemmli buffer (120mM Tris-HCl pH 7.5, 20% glycerol, 4% SDS, 0.02% bromophenol blue, 10% beta-mercaptoethanol) was added to each sample and incubated at 95°C for 5 min. Insoluble material was pelleted by centrifugation and an equal volume of the soluble fraction from each sample was run on an SDS-polyacrylamide gel (Mini-PROTEAN TGS Any kD precast gel; Bio-Rad, Hercules, CA Cat # 4569033) and transferred to a nitrocellulose membrane using a TransBlot Turbo Mini 0.2 μm Nitrocellulose Transfer Pack (Bio-Rad, Cat # 1704158) on the High MW setting of a TransBlot Turbo machine (Bio-Rad). The membrane was blocked in Intercept Blocking Buffer (LI-COR Biosciences, Lincoln, NE, Cat # 927-70001), and the following primary antibodies and dilutions were used for detection: V5 (R960-25, 1:5000; Invitrogen), Hxk2 (#100-4159, 1:20,000; Rockland Immunochemicals Inc., Pottstown, PA). The secondary antibodies used were IRDye 800CW (926-32210) and 680RD (926-68071) (1:20,000; LI-COR Biosciences), and the membrane was imaged on a LI-COR Odyssey Imager. All washing steps were performed with PBS + 0.1% Tween-20.

### Data Availability

All strains and plasmids are available upon request. ChIP-seq and DIP-seq data have been deposited in GEO under accession code GSE189038. Nanopore data was submitted to GEO and is currently being processed as a SuperSeries with GSE189038. This data will be available and publicly accessible at the time of publication.

## ACKNOWLEDGEMENTS

We would like to thank the Rine Lab for lively discussions and helpful comments through the development and execution of this project, especially Davis Goodnight. We also especially thank Marc Fouet for generous technical assistance with lab servers. We thank Koen Van den Berge and Sandrine Dudoit for advice on statistics and new ways of thinking about and approaching our Nanopore data. We thank Marcus Stoiber at Oxford Nanopore for help and advice with Megalodon. This research used the Savio computational cluster resource provided by the Berkeley Research Computing program at the University of California, Berkeley (supported by the UC Berkeley Chancellor, Vice Chancellor for Research, and Chief Information Officer). This work was funded by grants from the National Institutes of Health to JR (R35 GM139488). MB received support from a National Science Foundation Graduate Research Fellowship (Grant No. 1752814).

## REFERENCES

Abdulhay NJ, McNally CP, Hsieh LJ, Kasinathan S, Keith A, Estes LS, Karimzadeh M, Underwood JG, Goodarzi H, Narlikar GJ, Ramani V. 2020. Massively multiplex single-molecule oligonucleosome footprinting. Elife 9. doi:10.7554/eLife.59404

Altaf M, Utley RT, Lacoste N, Tan S, Briggs SD, Côté J. 2007. Interplay of chromatin modifiers on a short basic patch of histone H4 tail defines the boundary of telomeric heterochromatin. Mol Cell 28:1002–1014. doi:10.1016/j.molcel.2007.12.002

Altemose N, Maslan A, Smith OK, Sundararajan K, Brown RR, Detweiler AM, Neff N, Miga KH, Straight AF, Streets A. 2021. DiMeLo-seq: a long-read, single-molecule method for mapping protein-DNA interactions genome-wide. bioRxiv. doi:10.1101/2021.07.06.451383

Armache K-J, Garlick JD, Canzio D, Narlikar GJ, Kingston RE. 2011. Structural basis of silencing: Sir3 BAH domain in complex with a nucleosome at 3.0 Å resolution. Science 334:977–982. doi:10.1126/science.1210915

Babiarz JE, Halley JE, Rine J. 2006. Telomeric heterochromatin boundaries require NuA4-dependent acetylation of histone variant H2A.Z in Saccharomyces cerevisiae. Genes Dev 20:700–710. doi:10.1101/gad.1386306

Behrouzi R, Lu C, Currie MA, Jih G, Iglesias N, Moazed D. 2016. Heterochromatin assembly by interrupted Sir3 bridges across neighboring nucleosomes. Elife 5. doi:10.7554/eLife.17556

Braunstein M, Rose AB, Holmes SG, Allis CD, Broach JR. 1993. Transcriptional silencing in yeast is associated with reduced nucleosome acetylation. Genes Dev 7:592–604. doi:10.1101/gad.7.4.592

Brothers M, Rine J. 2019. Mutations in the PCNA DNA Polymerase Clamp of Saccharomyces cerevisiae Reveal Complexities of the Cell Cycle and Ploidy on Heterochromatin Assembly. Genetics 213:449–463. doi:10.1534/genetics.119.302452

Buchberger JR, Onishi M, Li G, Seebacher J, Rudner AD, Gygi SP, Moazed D. 2008. Sir3-Nucleosome Interactions in Spreading of Silent Chromatin in Saccharomyces cerevisiae. Mech Chem Biosyst 28:6903–6918. doi:10.1128/MCB.01210-08

Buchman AR, Kimmerly WJ, Rine J, Kornberg RD. 1988. Two DNA-binding factors recognize specific sequences at silencers, upstream activating sequences, autonomously replicating sequences, and telomeres in Saccharomyces cerevisiae. Mol Cell Biol 8:210–225. doi:10.1128/mcb.8.1.210

Carmen AA, Milne L, Grunstein M. 2002. Acetylation of the yeast histone H4 N terminus regulates its binding to heterochromatin protein SIR3. J Biol Chem 277:4778–4781. doi:10.1074/jbc.M110532200

Chang J-F, Hall BE, Tanny JC, Moazed D, Filman D, Ellenberger T. 2003. Structure of the coiled-coil dimerization motif of Sir4 and its interaction with Sir3. Structure 11:637–649. doi:10.1016/s0969-2126(03)00093-5

Chen L, Widom J. 2005. Mechanism of transcriptional silencing in yeast. Cell 120:37–48. doi:10.1016/j.cell.2004.11.030

Chereji RV, Ramachandran S, Bryson TD, Henikoff S. 2018. Precise genome-wide mapping of single nucleosomes and linkers in vivo. Genome Biol 19:19. doi:10.1186/s13059-018-1398-0

Cockell M, Palladino F, Laroche T, Kyrion G, Liu C, Lustig AJ, Gasser SM. 1995. The carboxy termini of Sir4 and Rap1 affect Sir3 localization: evidence for a multicomponent complex required for yeast telomeric silencing. J Cell Biol 129:909–924. doi:10.1083/jcb.129.4.909

Dodson AE, Rine J. 2015. Heritable capture of heterochromatin dynamics in Saccharomyces cerevisiae. Elife 4:e05007. doi:10.7554/eLife.05007

Donze D, Kamakaka RT. 2001. RNA polymerase III and RNA polymerase II promoter complexes are heterochromatin barriers in Saccharomyces cerevisiae. EMBO J 20:520–531. doi:10.1093/emboj/20.3.520

Ehrentraut S, Hassler M, Oppikofer M, Kueng S, Weber JM, Mueller JW, Gasser SM, Ladurner AG, Ehrenhofer-Murray AE. 2011. Structural basis for the role of the Sir3 AAA+ domain in silencing: interaction with Sir4 and unmethylated histone H3K79. Genes Dev 25:1835–1846. doi:10.1101/gad.17175111

Ellahi A, Thurtle DM, Rine J. 2015. The Chromatin and Transcriptional Landscape of Native Saccharomyces cerevisiae Telomeres and Subtelomeric Domains. Genetics 200:505–521. doi:10.1534/genetics.115.175711

Engreitz JM, Pandya-Jones A, McDonel P, Shishkin A, Sirokman K, Surka C, Kadri S, Xing J, Goren A, Lander ES, Plath K, Guttman M. 2013. The Xist lncRNA exploits three-dimensional genome architecture to spread across the X chromosome. Science 341:1237973. doi:10.1126/science.1237973

Galupa R, Heard E. 2018. X-Chromosome Inactivation: A Crossroads Between Chromosome Architecture and Gene Regulation. Annu Rev Genet 52:535–566. doi:10.1146/annurev-genet-120116-024611

Gao L, Gross DS. 2008. Sir2 Silences Gene Transcription by Targeting the Transition between RNA Polymerase II Initiation and Elongation. Mech Chem Biosyst 28:3979–3994. doi:10.1128/MCB.00019-08

Gartenberg MR, Smith JS. 2016. The Nuts and Bolts of Transcriptionally Silent Chromatin in Saccharomyces cerevisiae. Genetics 203:1563–1599. doi:10.1534/genetics.112.145243

Georgel PT, Palacios DeBeer MA, Pietz G, Fox CA, Hansen JC. 2001. Sir3-dependent assembly of supramolecular chromatin structures in vitro. Proc Natl Acad Sci U S A 98:8584–8589. doi:10.1073/pnas.151258798

Ghidelli S, Donze D, Dhillon N, Kamakaka RT. 2001. Sir2p exists in two nucleosome-binding complexes with distinct deacetylase activities. EMBO J 20:4522–4535. doi:10.1093/emboj/20.16.4522

Giaimo BD, Ferrante F, Herchenröther A, Hake SB, Borggrefe T. 2019. The histone variant H2A.Z in gene regulation. Epigenetics Chromatin 12:37. doi:10.1186/s13072-019-0274-9

Goodnight D, Rine J. 2020. S-phase-independent silencing establishment in Saccharomyces cerevisiae. Elife 9. doi:10.7554/eLife.58910

Gotta M, Palladino F, Gasser SM. 1998. Functional characterization of the N terminus of Sir3p. Mol Cell Biol 18:6110–6120. doi:10.1128/MCB.18.10.6110

Gottschling DE. 1992. Telomere-proximal DNA in Saccharomyces cerevisiae is refractory to methyltransferase activity in vivo.pdf. Proc Natl Acad Sci U S A 89:4062–4065.

Gottschling DE, Aparicio OM, Billington BL, Zakian VA. 1990. Position effect at S. cerevisiae telomeres: reversible repression of Pol II transcription. Cell 63:751–762. doi:10.1016/0092-8674(90)90141-z

Hecht A, Laroche T, Strahl-Bolsinger S, Gasser SM, Grunstein M. 1995. Histone H3 and H4 N-termini interact with SIR3 and SIR4 proteins: a molecular model for the formation of heterochromatin in yeast. Cell 80:583–592. doi:10.1016/0092-8674(95)90512-x

Hecht A, Strahl-Bolsinger S, Grunstein M. 1996. Spreading of transcriptional repressor SIR3 from telomeric heterochromatin. Nature 383:92–96.

Henikoff S. 1996. Dosage-dependent modification of position-effect variegation in Drosophila. Bioessays 18:401–409.

Hentges P, Van Driessche B, Tafforeau L, Vandenhaute J, Carr AM. 2005. Three novel antibiotic marker cassettes for gene disruption and marker switching in Schizosaccharomyces pombe. Yeast 22:1013–1019. doi:10.1002/yea.1291

Hocher A, Ruault M, Kaferle P, Descrimes M, Garnier M, Morillon A, Taddei A. 2018. Expanding heterochromatin reveals discrete subtelomeric domains delimited by chromatin landscape transitions. Genome Res 28:1867–1881. doi:10.1101/gr.236554.118

Hoppe GJ, Tanny JC, Rudner AD, Gerber SA, Danaie S, Gygi SP, Moazed D. 2002. Steps in assembly of silent chromatin in yeast: Sir3-independent binding of a Sir2/Sir4 complex to silencers and role for Sir2-dependent deacetylation. Mol Cell Biol 22:4167–4180. doi:10.1128/mcb.22.12.4167-4180.2002

Imai S, Armstrong CM, Kaeberlein M, Guarente L. 2000. Transcriptional silencing and longevity protein Sir2 is an NAD-dependent histone deacetylase. Nature 403:795–800. doi:10.1038/35001622

Johnson A, Li G, Sikorski TW, Buratowski S, Woodcock CL, Moazed D. 2009. Reconstitution of heterochromatin-dependent transcriptional gene silencing. Mol Cell 35:769–781. doi:10.1016/j.molcel.2009.07.030

Johnson A, Wu R, Peetz M, Gygi SP, Moazed D. 2013. Heterochromatic gene silencing by activator interference and a transcription elongation barrier. J Biol Chem 288:28771–28782. doi:10.1074/jbc.M113.460071

Johnson LM, Kayne PS, Kahn ES, Grunstein M. 1990. Genetic evidence for an interaction between SIR3 and histone H4 in the repression of the silent mating loci in Saccharomyces cerevisiae. Proc Natl Acad Sci USA 87:6286–6290.

Kimmerly W, Buchman A, Kornberg R, Rine J. 1988. Roles of two DNA-binding factors in replication, segregation and transcriptional repression mediated by a yeast silencer. EMBO J 7:2241–2253. doi:10.1002/j.1460-2075.1988.tb03064.x

King DA, Hall BE, Iwamoto MA, Win KZ, Chang JF, Ellenberger T. 2006. Domain structure and protein interactions of the silent information regulator Sir3 revealed by screening a nested deletion library of protein fragments. J Biol Chem 281:20107–20119. doi:10.1074/jbc.M512588200

Landry J, Sutton A, Tafrov ST, Heller RC, Stebbins J, Pillus L, Sternglanz R. 2000. The silencing protein SIR2 and its homologs are NAD-dependent protein deacetylases. Proc Natl Acad Sci U S A 97:5807–5811. doi:10.1073/pnas.110148297

Langmead B, Salzberg SL. 2012. Fast gapped-read alignment with Bowtie 2. Nat Methods 9:357–359. doi:10.1038/nmeth.1923

Lee ME, DeLoache WC, Cervantes B, Dueber JE. 2015. A Highly Characterized Yeast Toolkit for Modular, Multipart Assembly. ACS Synth Biol 4:975–986. doi:10.1021/sb500366v

Liaw H, Lustig AJ. 2006. Sir3 C-terminal domain involvement in the initiation and spreading of heterochromatin. Mol Cell Biol 26:7616–7631. doi:10.1128/MCB.01082-06

Locke J, Kotarski MA, Tartof KD. 1988. Dosage-dependent modifiers of position effect variegation in Drosophila and a mass action model that explains their effect. Genetics 120:181–198.

Longtine MS, Wilson NM, Petracek ME, Berman J. 1989. A yeast telomere binding activity binds to two related telomere sequence motifs and is indistinguishable from RAP1. Curr Genet 16:225–239. doi:10.1007/BF00422108

Loo S, Rine J. 1994. Silencers and Domains of Generalized Repression. Science 264:1748–1771.

Lynch PJ, Rusche LN. 2009. A Silencer Promotes the Assembly of Silenced Chromatin Independently of Recruitment. Mech Chem Biosyst 29:43–56. doi:10.1128/MCB.00983-08

Meneghini MD, Wu M, Madhani HD. 2003. Conserved histone variant H2A.Z protects euchromatin from the ectopic spread of silent heterochromatin. Cell 112:725–736. doi:10.1016/s0092-8674(03)00123-5

Moretti P, Freeman K, Coodly L, Shore D. 1994. Evidence that a complex of SIR proteins interacts with the silencer and telomere-binding protein RAP1. Genes Dev 8:2257–2269. doi:10.1101/gad.8.19.2257

Moretti P, Shore D. 2001. Multiple interactions in Sir protein recruitment by Rap1p at silencers and telomeres in yeast. Mol Cell Biol 21:8082–8094. doi:10.1128/MCB.21.23.8082-8094.2001

Murray IA, Morgan RD, Luyten Y, Fomenkov A, Corrêa IR Jr, Dai N, Allaw MB, Zhang X, Cheng X, Roberts RJ. 2018. The non-specific adenine DNA methyltransferase M.EcoGII. Nucleic Acids Res 46:840–848. doi:10.1093/nar/gkx1191

Ng HH, Ciccone DN, Morshead KB, Oettinger MA, Struhl K. 2003. Lysine-79 of histone H3 is hypomethylated at silenced loci in yeast and mammalian cells: a potential mechanism for position-effect variegation. Proc Natl Acad Sci U S A 100:1820–1825. doi:10.1073/pnas.0437846100

Ng HH, Feng Q, Wang H, Erdjument-Bromage H, Tempst P, Zhang Y, Struhl K. 2002. Lysine methylation within the globular domain of histone H3 by Dot1 is important for telomeric silencing and Sir protein association. Genes Dev 16:1518–1527. doi:10.1101/gad.1001502

Norris A, Bianchet MA, Boeke JD. 2008. Compensatory interactions between Sir3p and the nucleosomal LRS surface imply their direct interaction. PLoS Genet 4:e1000301. doi:10.1371/journal.pgen.1000301

Oki M, Valenzuela L, Chiba T, Ito T, Kamakaka RT. 2004. Barrier proteins remodel and modify chromatin to restrict silenced domains. Mol Cell Biol 24:1956–1967. doi:10.1128/mcb.24.5.1956-1967.2004

Onishi M, Liou G-G, Buchberger JR, Walz T, Moazed D. 2007. Role of the conserved Sir3-BAH domain in nucleosome binding and silent chromatin assembly. Mol Cell 28:1015–1028. doi:10.1016/j.molcel.2007.12.004

Oppikofer M, Kueng S, Keusch JJ, Hassler M, Ladurner AG, Gut H, Gasser SM. 2013. Dimerization of Sir3 via its C-terminal winged helix domain is essential for yeast heterochromatin formation. EMBO J 32:437–449. doi:10.1038/emboj.2012.343

Oppikofer M, Kueng S, Martino F, Soeroes S, Hancock SM, Chin JW, Fischle W, Gasser SM. 2011. A dual role of H4K16 acetylation in the establishment of yeast silent chromatin. EMBO J 30:2610–2621. doi:10.1038/emboj.2011.170

Park J-H, Cosgrove MS, Youngman E, Wolberger C, Boeke JD. 2002. A core nucleosome surface crucial for transcriptional silencing. Nat Genet 32:273–279. doi:10.1038/ng982

Radman-Livaja M, Ruben G, Weiner A, Friedman N, Kamakaka R, Rando OJ. 2011. Dynamics of Sir3 spreading in budding yeast: secondary recruitment sites and euchromatic localization. EMBO J 30:1012–1026. doi:10.1038/emboj.2011.30

Renauld H, Aparicio OM, Zierath PD, Billington BL, Chhablani SK, Gottschling DE. 1993. Silent domains are assembled continuously from the telomere and are defined by promoter distance and strength, and by SIR3 dosage. Genes & Development 7:1133–1145.

Rudner AD, Hall BE, Ellenberger T, Moazed D. 2005. A nonhistone protein-protein interaction required for assembly of the SIR complex and silent chromatin. Mol Cell Biol 25:4514–4528. doi:10.1128/MCB.25.11.4514-4528.2005

Rusche LN, Kirchmaier AL, Rine J. 2002. Ordered Nucleation and Spreading of Silenced Chromatin in Saccharomyces cerevisiae. Molecular Biology of the Cell 13:2207–2222. doi:10.1091/mbc.E02

Samel A, Rudner A, Ehrenhofer-Murray AE. 2017. Variants of the Sir4 Coiled-Coil Domain Improve Binding to Sir3 for Heterochromatin Formation in Saccharomyces cerevisiae. G3 7:1117–1126. doi:10.1534/g3.116.037739

Sampath V, Yuan P, Wang IX, Prugar E, van Leeuwen F, Sternglanz R. 2009. Mutational analysis of the Sir3 BAH domain reveals multiple points of interaction with nucleosomes. Mol Cell Biol 29:2532–2545. doi:10.1128/MCB.01682-08

Saxton DS, Rine J. 2019. Epigenetic memory independent of symmetric histone inheritance. Elife 8. doi:10.7554/eLife.51421

Schindelin J, Arganda-Carreras I, Frise E, Kaynig V, Longair M, Pietzsch T, Preibisch S, Rueden C, Saalfeld S, Schmid B, Tinevez J-Y, White DJ, Hartenstein V, Eliceiri K, Tomancak P, Cardona A. 2012. Fiji: an open-source platform for biological-image analysis. Nat Methods 9:676–682. doi:10.1038/nmeth.2019

Schnell R, Rine J. 1986. A position effect on the expression of a tRNA gene mediated by the SIR genes in Saccharomyces cerevisiae. Mol Cell Biol 6:494–501. doi:10.1128/mcb.6.2.494-501.1986

Sekinger EA, Gross DS. 2001. Silenced chromatin is permissive to activator binding and PIC recruitment. Cell 105:403–414. doi:10.1016/s0092-8674(01)00329-4

Shipony Z, Marinov GK, Swaffer MP, Sinnott-Armstrong NA, Skotheim JM, Kundaje A, Greenleaf WJ. 2020. Long-range single-molecule mapping of chromatin accessibility in eukaryotes. Nat Methods 17:319–327. doi:10.1038/s41592-019-0730-2

Shore D, Nasmyth K. 1987. Purification and cloning of a DNA binding protein from yeast that binds to both silencer and activator elements. Cell 51:721–732. doi:10.1016/0092-8674(87)90095-x

Shore D, Stillman DJ, Brand AH, Nasmyth KA. 1987. Identification of silencer binding proteins from yeast: possible roles in SIR control and DNA replication. EMBO J 6:461–467. doi:10.1002/j.1460-2075.1987.tb04776.x

Simms TA, Dugas SL, Gremillion JC, Ibos ME, Dandurand MN, Toliver TT, Edwards DJ, Donze D. 2008. TFIIIC binding sites function as both heterochromatin barriers and chromatin insulators in Saccharomyces cerevisiae. Eukaryot Cell 7:2078–2086. doi:10.1128/EC.00128-08

Simon MD, Pinter SF, Fang R, Sarma K, Rutenberg-Schoenberg M, Bowman SK, Kesner BA, Maier VK, Kingston RE, Lee JT. 2013. High-resolution Xist binding maps reveal two-step spreading during X-chromosome inactivation. Nature 504:465–469. doi:10.1038/nature12719

Singh J, Klar AJ. 1992. Active genes in budding yeast display enhanced in vivo accessibility to foreign DNA methylases: a novel in vivo probe for chromatin structure of yeast. Genes Dev 6:186–196. doi:10.1101/gad.6.2.186

Smith JS, Brachmann CB, Celic I, Kenna MA, Muhammad S, Starai VJ, Avalos JL, Escalante-Semerena JC, Grubmeyer C, Wolberger C, Boeke JD. 2000. A phylogenetically conserved NAD+-dependent protein deacetylase activity in the Sir2 protein family. Proc Natl Acad Sci U S A 97:6658–6663. doi:10.1073/pnas.97.12.6658

Stavenhagen JB, Zakian VA. 1994. Internal tracts of telomeric DNA act as silencers in Saccharomyces cerevisiae. Genes Dev 8:1411–1422. doi:10.1101/gad.8.12.1411

Steakley DL, Rine J. 2015. On the Mechanism of Gene Silencing in Saccharomyces cerevisiae. G3: Genes, Genomes, Genetics 5:1751–1763. doi:10.1534/g3.115.018515

Stergachis AB, Debo BM, Haugen E, Churchman LS, Stamatoyannopoulos JA. 2020. Single-molecule regulatory architectures captured by chromatin fiber sequencing. Science 368:1449–1454. doi:10.1126/science.aaz1646

Stone EM, Reifsnyder C, McVey M, Gazo B, Pillus L. 2000. Two classes of sir3 mutants enhance the sir1 mutant mating defect and abolish telomeric silencing in Saccharomyces cerevisiae. Genetics 155:509–522.

Strahl-Bolsinger S, Hecht A, Luo K, Grunstein M. 1997. SIR2 and SIR4 interactions differ in core and extended telomeric heterochromatin in yeast. Genes Dev 11:83–93. doi:10.1101/gad.11.1.83

Stulemeijer IJ, Pike BL, Faber AW, Verzijlbergen KF, van Welsem T, Frederiks F, Lenstra TL, Holstege FC, Gasser SM, van Leeuwen F. 2011. Dot1 binding induces chromatin rearrangements by histone methylation-dependent and -independent mechanisms. Epigenetics Chromatin 4:2. doi:10.1186/1756-8935-4-2

Suka N, Suka Y, Carmen AA, Wu J, Grunstein M. 2001. Highly Specific Antibodies Determine Histone Acetylation Site Usage in Yeast Heterochromatin and Euchromatin. Mol Cell 8:473–479.

Sussel L, Vannier D, Shore D. 1993. Epigenetic switching of transcriptional states: cis- and trans- acting factors affecting establishment of silencing at the HMR locus in Saccharomyces cereivisae. Molecular and Cellular Biology 13:3919–3928.

Swygert SG, Senapati S, Bolukbasi MF, Wolfe SA, Lindsay S, Peterson CL. 2018. SIR proteins create compact heterochromatin fibers. Proc Natl Acad Sci U S A 115:12447–12452. doi:10.1073/pnas.1810647115

Thorvaldsdóttir H, Robinson JT, Mesirov JP. 2013. Integrative Genomics Viewer (IGV): high-performance genomics data visualization and exploration. Brief Bioinform 14:178–192. doi:10.1093/bib/bbs017

Thurtle DM, Rine J. 2014. The molecular topography of silenced chromatin in Saccharomyces cerevisiae. Genes Dev 28:245–258. doi:10.1101/gad.230532.113

Triolo T, Sternglanz R. 1996. Role of interactions between the origin recognition complex and SIR1 in transcriptional silencing. Nature 381:251–253.

Valenzuela L, Dhillon N, Kamakaka RT. 2009. Transcription independent insulation at TFIIIC-dependent insulators. Genetics 183:131–148. doi:10.1534/genetics.109.106203

Venkatasubrahmanyam S, Hwang WW, Meneghini MD, Tong AHY, Madhani HD. 2007. Genome-wide, as opposed to local, antisilencing is mediated redundantly by the euchromatic factors Set1 and H2A.Z. Proc Natl Acad Sci U S A 104:16609–16614. doi:10.1073/pnas.0700914104

Woodcock CB, Horton JR, Zhang X, Blumenthal RM, Cheng X. 2020. Beta class amino methyltransferases from bacteria to humans: evolution and structural consequences. Nucleic Acids Res. doi:10.1093/nar/gkaa446

Xu L, Seki M. 2020. Recent advances in the detection of base modifications using the Nanopore sequencer. J Hum Genet 65:25–33. doi:10.1038/s10038-019-0679-0

